# Postnatal SETD1B is essential for learning and the regulation of neuronal plasticity genes

**DOI:** 10.1101/2021.08.02.454636

**Authors:** Alexandra Michurina, M Sadman Sakib, Cemil Kerimoglu, Dennis Manfred Krüger, Lalit Kaurani, Md Rezaul Islam, Joshi Parth Devesh, Sophie Schröder, Tonatiuh Pena Centeno, Jiayin Zhou, Ranjit Pradhan, Julia Cha, Xingbo Xu, Gregor Eichele, Elisabeth M. Zeisberg, Andrea Kranz, A. Francis Stewart, Andre Fischer

## Abstract

Histone 3 lysine 4 methylation (H3K4me) is mediated by six different lysine methyltransferases. Amongst these enzymes SET domain containing 1b (SETD1B) has been linked to syndromic intellectual disability but its role in the postnatal brain has not been studied yet. Here we employ mice that lack *Setd1b* from excitatory neurons of the postnatal forebrain and combine neuron-specific ChIP-seq and RNA-seq approaches to elucidate its role in neuronal gene expression. We observe that SETD1B controls the expression of genes with a broad H3K4me3 peak at their promoters that represent neuronal enriched genes linked to learning and memory function. Comparative analysis to corresponding data from conditional *Kmt2a* and *Kmt2b* knockout mice suggests that this function is specific to SETD1B. Moreover, postnatal loss of *Setd1b* leads to severe learning impairment, suggesting that SETD1B-mediated regulation of H3K4me levels in postnatal neurons is critical for cognitive function.

## INTRODUCTION

Adult-onset cognitive diseases, such as for example sporadic Alzheimer’s disease (AD), depend on complex interactions of genetic and environmental risk factors that activate epigenetic processes (Fischer, 2014). In addition, mutations in genes that control epigenetic gene regulation are over-represented in neurodevelopmental diseases linked to cognitive dysfunction (Kleefstra *et al*, 2014) (Gabriele *et al*, 2018). Therefore, targeting the epigenome has emerged as a promising therapeutic avenue to treat neurodevelopmental and adult-onset cognitive diseases (Fischer, 2014; Nestler *et al*, 2015) (Larizza & Finelli, 2019). To understand the regulation of epigenetic gene expression in in the context of cognitive function is thus of utmost importance. Increasing evidence hints to a especially important role histone 3 lysine 4 (H3K4me) methylation, a mark linked to active transcription. In mammals, H3K4 methylation is mediated by six different lysine methyltransferases (KMTs), namely KMT2A (Mll1), KMT2B (Mll2), KMT2C (Mll3), KMT2D (Mll4), SETD1A, and SETD1B that catalyze – with some specificity - mono-, di- and trimethylation (Shilatifard, 2012). Importantly, mutations in all of these enzymes are found in neurodevelopmental intellectual disability disorders and a substantial amount of work has focused on the role of these enzymes in cortical development (Gabriele *et al.*, 2018). Neuronal H3K4me is also deregulated in adult-onset neurodegenerative diseases, such as Alzheimer’s disease (AD) (Gjoneska *et al*, 2015). Moreover, exposure of mice to a learning task was shown to increase H3K4me levels in postnatal brain (Gupta *et al*, 2010). However, the role of the different H3K4 KMTs in the postnatal brain is not well understood yet. Thus, it is presently also unclear to what extent postnatal processes may contribute to the cognitive phenotypes observed in intellectual disability disorders that are linked to mutations in the different H3K4 KMTs. Such knowledge is important for the development of therapeutic approaches. Amongst the six H3K4 KMTs, SETD1B is the least studied enzyme. Virtually nothing is known about the function of SETD1B in postnatal brain, although mutations in the *Setd1b* gene have been linked to syndromic intellectual disability (Hiraide *et al*, 2018) (Labonne *et al*, 2016) (Roston *et al*, 2020). Therefore, to elucidate the role of *Setd1b* we generated mice that lack the gene from excitatory neurons of the postnatal brain and analyzed cognitive function and epigenetic gene expression in young adult mice. Our data reveal that *Setd1b* is essential for memory formation. Moreover, we provide evidence that *Setd1b* controls the expression of genes that are characterized by a broad H3K4 trimethylation peak around the transcription start site (TSS) and are intimately linked to neuronal function and learning behavior. A comparative analysis to corresponding data from conditional *Kmt2a* and *Kmt2b* knockout mice suggest a distinct role for *Setd1b* in the regulation of epigenetic gene expression linked to learning processes.

## RESULTS

### Loss of Setd1b in postnatal forebrain neurons impairs hippocampus-dependent memory formation

To study the role of *Setd1b* in the postnatal brain, we crossed mice in which exon 5 of the *Setd1b* gene is flanked by loxP sites to mice that express CRE-recombinase under control of the CamKII promoter (Minichiello *et al*, 1999). This approach ensures deletion of *Setd1b* from excitatory forebrain neurons of the postnatal brain (cKO mice). Quantitative PCR (qPCR) analysis confirmed decreased expression of *Setd1b* in the hippocampal Cornu Ammonis (CA) area, the dentate gyrus (DG) and the cortex when compared to corresponding control littermates that carry loxP sites but do not express CRE recombinase (control group). Expression in the cerebellum was not affected, confirming the specificity of the approach (**Fig 1A**). Residual expression of *Setd1b* in the forebrain regions is most likely due to the fact that deletion is restricted to excitatory neurons while other cell types are unaffected. In line with the qPCR data, SETD1B protein levels were reduced in the hippocampal CA region of *Setd1b* cKO mice (**Fig 1B**). *Setd1b* cKO mice did not show any gross abnormalities in brain anatomy as evidenced by immunohistological analysis of DAPI staining, staining of marker proteins for neuronal integrity Neuronal N (NeuN), microtubule-associated protein 2 (MAP2) as well as ionized calcium-binding adapter molecule 1 (IBA1) as a marker for microglia and glial fibrillary acidic protein (GFAP) as a marker for astrocytes (**Fig. 1C**). Next, we subjected *Setd1b* cKO and control mice to behavior testing. Notably, it was previously shown that heterozygous mice expressing CRE under control of the CamKII promoter do not differ from wild type littermates (Kuczera *et al*, 2010) (Stilling *et al*, 2014)and we have confirmed this in the context of the present study also for behavior testing (**Fig. EV1**). There was no difference amongst groups in the open field test, suggesting that explorative behavior and basal anxiety is normal in *Setd1b* cKO mice (**Fig 1D**). Short term memory was assayed via the Y-maze and was also similar amongst groups (**Fig 1E**). We also tested depressive-like behavior in the Porsolt forced swim test. No difference was observed amongst groups (**Fig 1F**). Next, we subjected mice to the Morris Water Maze test to study hippocampus-dependent spatial memory. While control mice were able to learn the task as indicated by a reduced escape latency throughout the 10 days of training, *Setd1b* cKO mice were severely impaired (**Fig 1G**). We also performed a more sensitive analysis using a modified version of the MUST-C algorithm to measure the different spatial strategies that represent either hippocampus-dependent or independent abilities (Illouz *et al*, 2016) (Islam *et al*, 2021). Our results show that *Setd1b* cKO mice fail to adopt hippocampus-dependent search strategies such as “direct”, “corrected” and “short-chaining” (**Fig 1H**). Consistently, the cumulative learning score calculated on the basis of these search strategies showed severe learning impairment in *Setd1b* cKO mice (**Fig 1I**). Since memory acquisition was severely impaired in *Setd1b* cKO mice, and we moreover observed a trend for reduced swim speed during a probe test that was performed after the end of the training procedure, memory recall could not be analyzed (**Fig. EV2**). In conclusion, our data show that deletion of *Setd1b* from excitatory neurons of the postnatal forebrain leads to severe impairment of hippocampus-dependent learning abilities.

**Figure 1.**
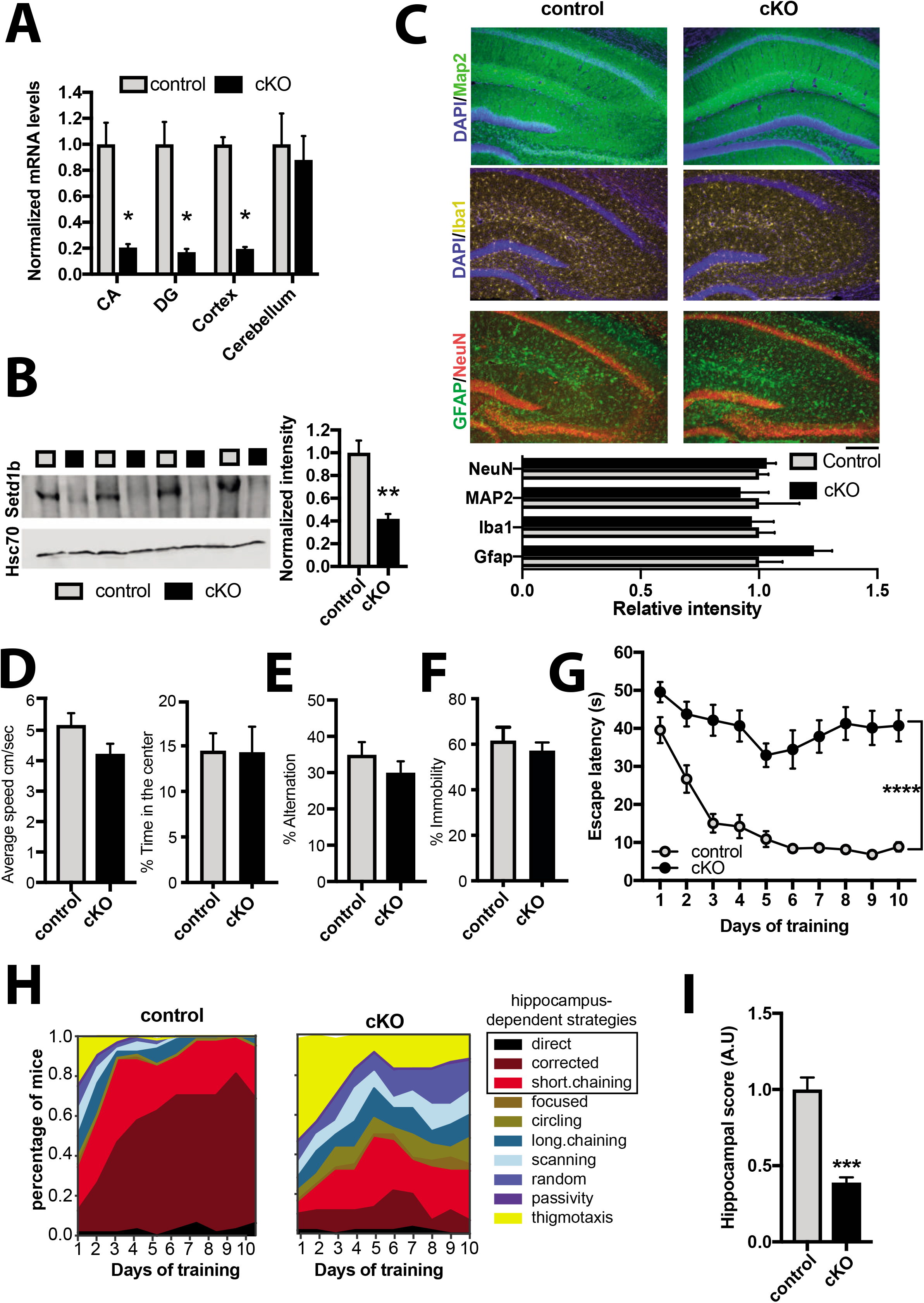
*Setd1b* is required for hippocampus-dependent memory. **A.** qPCR analysis shows loss of *Setd1b* in forebrain regions while levels in the cerebellum are not affected (CA: Control, n = 6; cKO, n = 6. DG: Control, n = 6; cKO, n = 6. Cortex: Control, n = 7; cKO, n = 7. Cerebellum: Control, n = 4; cKO, n = 4). * *p*-value < 0.05 (Student t-test). **B.** Immunoblot analysis shows loss of SETD1B protein in the hippocampus of *Setd1b* cKO mice (Control, n = 4; cKO, n = 4). ** *p*-value < 0.01 (Student t-test). **C.** Immunohistochemical staining (upper level) for marker proteins of neuronal integrity and quantification (lower panel) show no difference in control and *Setd1b* cKO mice (NeuN: Control, n = 6; cKO, n = 6; Student t-test *p*-value = 0.57. MAP2: Control, n = 4; cKO, n = 4; Student t-test *p*-value = 0.72. Iba1: Control, n = 5; cKO, n = 5; Student t-test *p*-value = 0.8. Gfap: Control, n = 5; cKO, n = 5; Student t-test *p*-value = 0.09.). Scale bar: 100 μm. **D.** Average speed (left panel) and percent time spent in the center (right panel) during exposure to the open field test was similar in control and *Setd1b* cKO mice (Average speed: Control, n = 15; cKO, n = 15; Student t-test *p*-value = 0.075. Time spent in center: Control, n = 15; cKO, n = 15; Student t-test *p*-value = 0.96). **E.** Short term memory was not different between control and *Setd1b* cKO mice as indicated by similar percent of alternations in the Y-maze test (Control, n = 15; cKO, n = 15; Student t-test *p*-value = 0.3). **F.** Depressive-like behavior was measured in the Porsolt forced swim test. There was no significant difference in the % time spent immobile between groups (Control, n = 8; cKO, n = 8; Student t-test *p*-value = 0.5). **G.** Escape latency during water maze training indicated severe learning impairment in *Setd1b* cKO mice (Control: n = 15, cKO: n = 15. Repeated measures ANOVA, genotype effect: F (1,28) = 82.34, **** *p*-value < 0.0001). **H.** Plots showing the specific search strategies during water maze training. Note the failure of *Setd1b* cKO mice to adopt hippocampus-dependent search strategies. **I.** The cognitive score calculated on the basis of the hippocampal search strategies reflects severe memory impairment in *Setd1b* cKO mice (Student t-test: *** p-value < 0.001).

### Setd1d controls H3K4 methylation peak width in hippocampal excitatory neurons

Although the Morris water maze test is used to specifically measure hippocampus-dependent learning and memory, we cannot exclude that other brain regions contribute to the observed learning impairment. Nevertheless, our data suggest that the hippocampus would be a suitable brain region to study the molecular function of neuronal *Setd1b* in the postnatal brain. We reasoned that analyzing the hippocampus would furthermore allow for an optimal comparison to previous studies that investigated epigenetic gene expression from hippocampal neurons and tissue obtained from *Kmt2a* and *Kmt2b* cKO mice, that were generated using the same CRE-driver line as in this study (Kerimoglu *et al*, 2013; Kerimoglu *et al*, 2017). To this end we isolated the hippocampal CA region from *Setd1b* cKO and control mice and prepared nuclei to perform neuron-specific chromatin immunoprecipitation (ChIP) (**Fig 2A**). Since SETD1B is a histone 3 lysine 4 (H3K4) methyltransferase we decided to analyze tri-methylation (H3K4me3) of histone 3 lysine 4 that is enriched at the transcription start site (TSS) of active genes and is associated with euchromatin and active gene expression. H3K4 methylation is believed to be a stepwise process and recent data suggest that the different methylation states (from mono- to tri-methylation) at the TSS of a gene form a gradient reflecting its specific transcriptional state (Choudhury *et al*, 2019; Soares *et al*, 2017). Moreover, some specificity for mono- di- or tri-methylation has been reported for the different H3K4 KMTs, although biochemical analysis does not always correspond to the *in vivo* data (Bochyńska *et al*, 2018; Kerimoglu *et al.*, 2017) (Gabriele *et al.*, 2018). Thus, we also analyzed mono-methylation of histone 3 at lysine 4 (H3K4me1). In addition, we analyzed histone 3 lysine 9 acetylation (H3K9ac), a euchromatin mark that was shown to partially depend on H3K4 methylation (Kerimoglu *et al.*, 2013). Finally, we also performed ChIP-Seq for histone 3 lysine 27 acetylation (H3K27ac), another euchromatin mark that is linked to active gene expression and marks promoter elements around the TSS but also enhancer regions and has not been directly linked to H3K4me3 in brain tissue. In all ChIP-Seq experiments we observed a clear separation between control and *Setd1b* cKO samples as evidenced by principal component analysis (**Fig EV3**).

**Figure 2.**
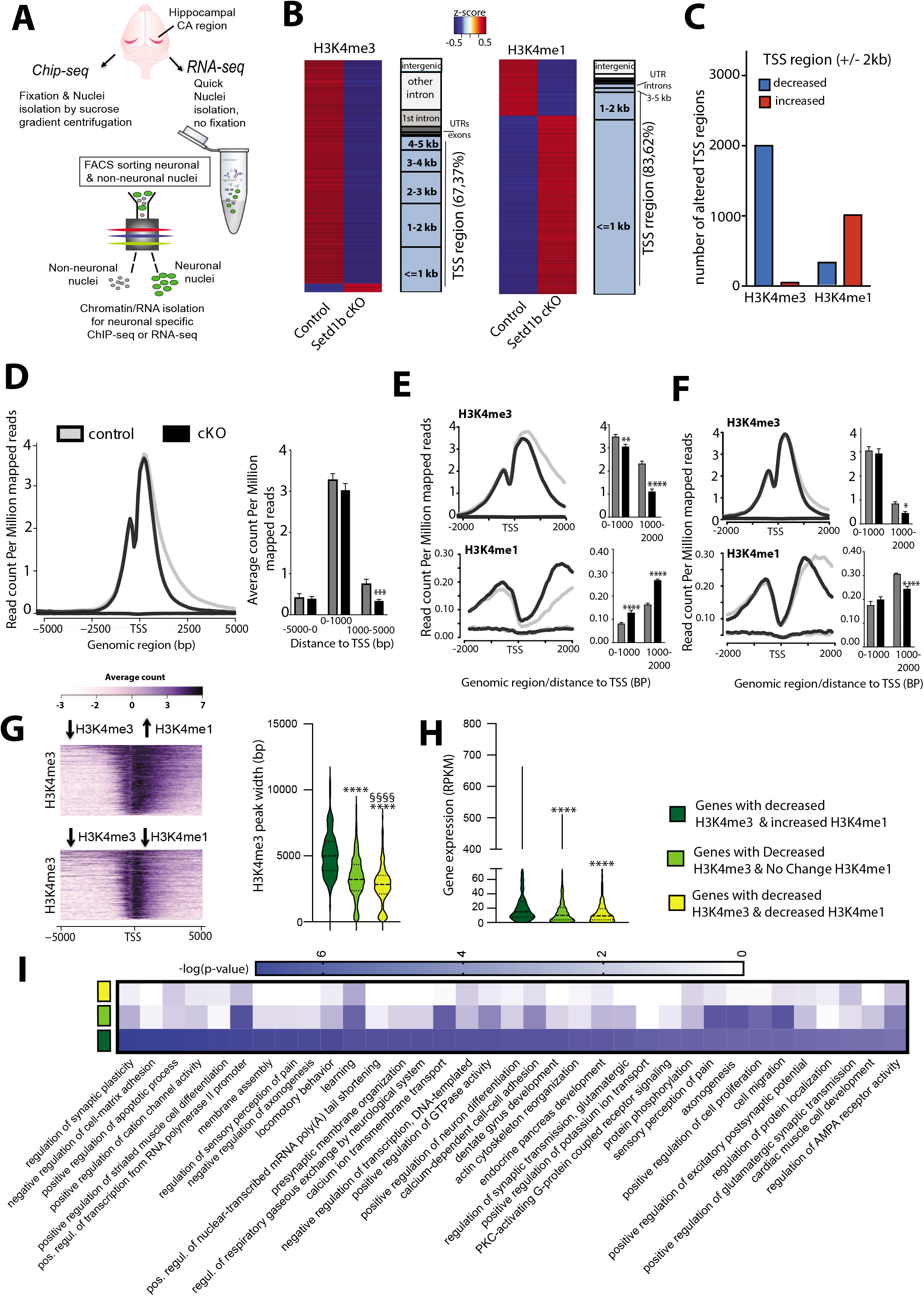
*Setd1b* controls H3K4 methylation and H3K4me3 peak width. **A.** Experimental scheme showing our approach to perform cell type specific ChIP-Seq and RNA-Seq. For ChIP-Seq we employed n = 4 for control and n = 4 for *Setd1b* cKO. **B.** Left panel: Heat map showing genomic regions with significantly differing H3K4me3 sites at the close vicinity of TSS (+/−2kb) in *Setd1b* cKO mice and the overall genomic locations of altered H3K4me3 levels. Right panel shows the same analysis for H3K4me1 (FDR < 0.05 & |fold change| > 1.5). **C.** Bar plot showing the number of TSS regions (+/− 2kb) with decreased and increased H3K4me3 and H3K4me1 in *Setd1b* cKO mice (FDR < 0.05 & |fold change| > 1.5). **D.** Profile plot showing H3K4me3 across all genes with significantly reduced H3K4me3 in *Setd1b* cKO mice. Left panel shows a bar chart indicating that reduced H3K4me3 in *Setd1b* cKO mice is mainly occurring downstream of the TSS (*** Student t-test *p*-value < 0.001). **E.** Profile plots showing the distribution of H3K4me3 and H3K4me1 at the close vicinity of TSS of genes that show significantly reduced H3K4me3 and increased H3K4me1 in *Setd1b* cKO mice. Bar graphs on the right show corresponding quantification (Student t-test: ** *p*-value < 0.01, **** *p*-value < 0.0001). **F.** Profile plots showing the distribution of H3K4me3 and H3K4me1 at the TSS of genes that show both reduced H3K4me3 and H3K4me1 in *Setd1b* cKO mice. Bar graphs on the left show corresponding quantification (Student t-test: * *p*-value < 0.05, **** *p*-value < 0.0001). **G.** Heatmap (left panel) showing basal state H3K4me3 width around TSS of genes characterized by decreased H3K4me3 in combination with either increased or decreased H3K4me1 in *Setd1b* cKO mice. Right panel: Quantification of the basal state H3K4me3 width around TSS of genes with decreased H3K4me3 in combination with either increased (472 peaks), decreased (401 peaks) or not altered H3K4me1 (1051 peaks) in *Setd1b* cKO mice (One-way ANOVA: *p*-value < 0.0001. Post-hoc multiple comparisons, Tukey’s test: increased H3K4me1 vs no change H3K4me1, **** *p*-value < 0.0001; increased H3K4me1 vs decreased H3K4me1, **** *p*-value < 0.0001; no change H3K4me1 vs decreased H3K4me1, §§§§ *p*-value < 0.0001). **H.** Violin plot showing the basal expression level for the 3 categories of genes that display decreased H3K4me3 in *Setd1b* cKO mice, now analyzed in wild type mice (increased H3K4me1: 456 genes, decreased H3K4me1: 382 genes, not altered H3K4me1: 993 genes). The basal expression level is highest for genes with decreased H3K4me3 in combination with increased H3K4me1 that are characterized by broad H3K4me3 peaks (One-way ANOVA: *p*-value < 0.0001. Post-hoc multiple comparisons, Tukey’s test: increased H3K4me1 vs no change H3K4me1, **** *p*-value < 0.0001; increased H3K4me1 vs decreased H3K4me1, **** *p*-value < 0.0001; no change H3K4me1 vs decreased H3K4me1, *p*-value = 0.6967). Please note that the number of genes is smaller than the number of peaks in (G) because in some cases there are more than one peaks around TSS of the same gene. **I.** Heat map showing functional pathways for the 3 categories of genes affected by reduced H3K4me3 in *Setd1b* cKO mice. Error bars indicate SEM.

Loss of Setd1b leads to a substantial decrease in neuronal H3K4me3 (adjusted *p*-value < 0.05 & fold change < −1.5) across the genome while the majority of significant changes are localized to regions in proximity to TSS (**Fig 2B, C; Supplementary Table 1**). Similar changes were observed for neuronal H3K9ac and H3K27ac, although fewer regions were affected when compared to H3K4me3 (**Fig. EV4; Supplementary Tables 2 and 3**). The changes in H3K9ac showed a substantial convergence with altered H3K4me3 (**Fig EV4C**). Thus, almost all TSS regions (+/− 2kb) exhibiting decreased H3K9ac were also marked by decreased H3K4me3, while the TSS regions showing decreased H3K27ac only partially overlapped with those with decreased H3K4me3 (**Fig EV4C**). These data support previous findings, showing that H3K4me3 is functionally linked to H3K9ac (Kerimoglu *et al.*, 2013) and suggest that the observed changes in H3K27ac might also reflect secondary effects.

We also observed significantly altered H3K4me1 levels (adjusted *p*-value < 0.05 & |fold change| < 1.5) in neurons of *Setd1b* cKO mice (**Fig 2B; Supplementary Table 4**). These changes were also almost exclusively detected in vicinity to the TSS (**Fig 2B**) but in contrast to the other investigated histone modifications, many of the significantly altered genomic regions exhibited increased H3K4me1 levels in *Setd1b* cKO mice (**Fig. 2B, C**), suggesting that *Setd1b* may mainly promotes tri-methylation of H3K4 in neurons. To further interpret these observations, we analyzed the distribution of altered H3K4 methylation across the TSS region including up- and downstream regions up to 5 kb. Interestingly, decreased H3K4me3 in cKO manifested itself exclusively downstream of the TSS, indicating that loss of *Sedt1b* may affect peak width (**Fig 2D**). We decided to further explore this observation and detected two distinct patterns of altered H3K4me. Namely, there was a difference between the genes that exhibit decreased H3K4me3 and at the same time increased H3K4me1 (**Fig. 2E**) and those that show concomitantly decreased H3K4 tri- and mono-methylation around the TSS (**Fig 2F**). The change in H3K4me3 was most significant in genes with decreased H3K4me3 and increased H3K4me1 and was characterized by a substantially reduced H3K4me3 peak width (**Fig 2E**), when compared to genes with milder yet significantly decreased H3K4me3 and H3K4me1 (**Fig 2F**). Findings from other cell types suggest a gradient of H3K4 methylation states in which the proximity of the mark to the TSS is correlated to the level of gene expression. Thus, genes with broader H3K4me3 peaks at the TSS exhibit the highest and most consistent expression levels and represent genes of particular importance for cellular identity and function (Benayoun *et al*, 2015) (Soares *et al.*, 2017). Our data revealed that the genes which are characterized by decreased H3K4me3 and increased H3K4me1 in *Setd1b* cKO mice, exhibit significantly broader H3K4me3 peaks under basal conditions, when compared to genes characterized by decreased H3K4me3 but either decreased or unchanged H3K4me1 levels (**Fig 2G; Supplementary Table 5**). Interestingly, these genes were also expressed at significantly higher levels under baseline conditions (**Fig 2H; Supplementary Table 6**). Gene-ontology (GO) analysis revealed that the genes with decreased H3K4me3 and increased H3K4me1 and thus having the broadest H3K4me3 peaks under basal conditions, represent pathways intimately linked to the function of excitatory hippocampal neurons (**Fig 2I; Supplementary Table 7**). Most importantly, this was not the case for the genes of the other two categories (**Fig 2I**). Finally, through motif enrichment analysis we observed that the TSS regions with decreased H3K4me3 and increased H3K4me1 show a significant enrichment for RE1 Silencing Transcription Factor (REST) consensus binding site (**Supplementary Table 8**), which is in line with our GO-term analysis since the REST complex has previously been shown to repress genes specifically important for synaptic plasticity and specific to neuronal function both during development and aging by recruiting an enzymatic machinery that mediates heterochromatin formation and thus acts as counterplayer the H3K4 methyltransferases such as SETD1B (Hwang & Zukin, 2018).

In summary, our data show that loss of Setd1b from hippocampal excitatory neurons leads to distinct changes in neuronal histone methylation and point to a specific role of SETD1B in the H3K4 tri-methylation and the expression of genes essential for hippocampal excitatory neuronal function.

### Setd1b controls the levels of highly expressed hippocampal excitatory neuronal genes characterized by a broad H3K4me3 peak at the TSS

To test the impact of SETD1B on gene expression directly, we analyzed RNA-sequencing data obtained from neuronal nuclei of the same hippocampi used to generate ChIP-seq data. This was possible since we had employed a modified fixation protocol that allowed us to perform neuron-specific ChIP-seq and RNA-seq from the same samples (**See Fig 2A, Fig EV5**). In line with the established role of H3K4me3 in active gene expression, we mainly detected downregulated genes when comparing control to *Setd1b* cKO mice (adjusted *p*-value < 0.1 & |fold change| < 1.2) (**Fig 3A, Fig EV6; Supplementary Table 9**). The comparatively few up-regulated genes mainly exhibited low expression at baseline (RPKM down-regulated genes =18.77 +/−1.45 vs. RPKM up-regulated genes = 3.08 +/− 0.26; Student’s t-test *p-value* < 0.0001). Further analysis revealed that the TSS region of genes down-regulated in *Setd1b* cKO mice is characterized by significantly reduced H3K4me3 peak width and increased H3K4me1 (**Fig 3B, 3D**), which was not the case for an equally sized random sample of unaffected genes (**Fig 3C**). H3K9ac and H3K27ac levels were also reduced at the TSS of the down-regulated genes (**Fig EV7**). This observation was specific to the genes down-regulated in *Setd1b* cKO mice, since random sets of genes that were not dysregulated in *Setd1b* cKO mice show normal H3K4me3, H3K4me1, H3K9ac and H3K27ac levels at the TSS (**Fig 3C, Fig EV7**). We also observed that the genes downregulated as a result of *Setd1b* deletion were characterized by a significantly broader H3K4me3 peak around TSS and higher expression under basal conditions, when compared to unaffected genes (**Fig 3E; Supplementary Tables 10 and 11**).

**Figure 3.**
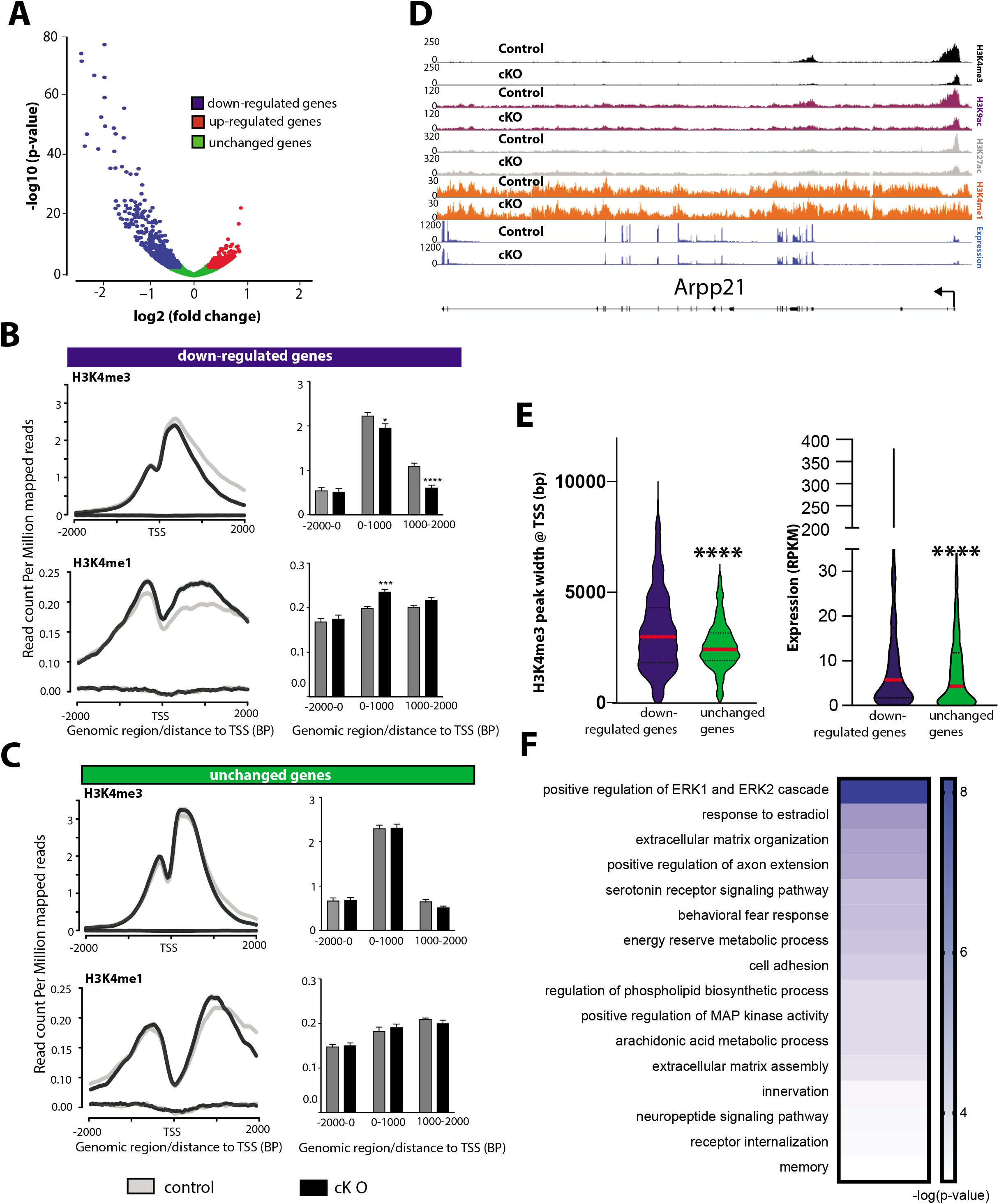
Hippocampal Setd1b controls highly expressed learning and memory genes characterized by a broad H3K4me3 peak. **A.** nucRNA-seq experiment. Volcano plot showing genes differentially expressed in hippocampal CA neurons of *Setd1b* cKO mice. n = 3/group. **B.** Profile plots showing H3K4me3 and H3K4me1 at the TSS of genes downregulated in *Setd1b* cKO mice. Bar plots (right panel) show quantification (Student t-test: * *p*-value < 0.05, *** *p*-value < 0.001, **** *p*-value < 0.0001). **C.** Profile plots showing H3K4me3 and H3K4me1 at the TSS of a random set of genes (equal in number to downregulated genes) that were not altered in *Setd1b* cKO mice. Bar plot (right panel) shows quantifications. **D.** *Arpp21* (CAMP Regulated Phosphoprotein 21) was selected as a representative gene downregulated in hippocampal neurons of *Setd1b* cKO mice to illustrate changes in the analyzed histone modifications. Please note that the H3K4me3 peak width is substantially shrinking in *Setd1b* cKO mice. At the same time there is an obvious increase of H3K4me1 at the TSS of *Arpp21* in *Setd1b* cKO mice. **E.** Left panel: H3K4me3 peaks are significantly broader in genes that are downregulated in *Setd1b* cKO mice, when compared to a random set of genes of equal number that were unaffected. Right panel: Genes downregulated in *Setd1b* cKO mice are characterized by higher baseline expression when compared to a random set of genes of equal number that were unaffected (Student t-test: **** *p*-value < 0.0001). **F.** Heat map showing functional pathways affected by genes downregulated in *Sed1b* cKO mice. Error bars indicate SEM.

GO analysis revealed that the genes down-regulated in *Setd1b* cKO mice are linked to synaptic plasticity and learning related processes such as for example ERK1/2 signaling, extracellular matrix organization or neuropeptide signaling pathways(Borbély *et al*, 2013; Sweatt, 2001) (Tsilibary *et al*, 2014) (**Fig 3F; Supplementary Table 12**). A specific example is the *Arpp-21* gene that leads to aberrant dendritic plasticity when down-regulated (Rehfeld *et al*, 2018) and has been genetically linked to intellectual disability (Marangi *et al*, 2013) or the *Npy2r* gene that leads to memory impairment in corresponding knockout mice (Redrobe *et al*) (**Fig EV8**). *Neuropsin* (*Klk8*) is another gene affected in *Setd1b* cKO mice that plays a complex role in memory function. It regulates the extracellular matrix, leads to spatial reference memory impairment when deleted in mice (Tamura *et al*, 2006) and has been implicated with mental diseases (Bukowski *et al*, 2020) (**Fig EV8**).

In summary, our data further suggest that SETD1B controls a specific set of genes that are characterized by a broad H3K4me3 peak at the TSS, are highly expressed in hippocampal excitatory neurons under basal conditions and play a specific role in synaptic plasticity.

### Comparison of epigenetic neuronal gene expression in *Kmt2a*, *Kmt2b* and *Setd1b* cKO mice

To further elucidate the role of *Setd1b* in the regulation of neuronal gene expression we decided to compare the data from *Setd1b* cKO mice to those from other mammalian H3K4 KMTs. We have previously generated comparable H3K4me3 and H3K4me1 ChIP-Seq data from neuronal nuclei obtained from the hippocampal CA region of conditional knockout mice that lack either *Kmt2a* or *Kmt2b* from excitatory forebrain neurons using the same CRE-line as employed in this study (Kerimoglu *et al.*, 2017). Since these experiments were performed at different timepoints we reanalyzed in parallel the H3K4me3 and H3K4me1 ChIP-Seq datasets obtained from hippocampal neuronal nuclei of *Kmt2a*, *Kmt2b* and *Setd1b* cKO mice (cut off adjusted p-value < 0.1 & |fold change| > 1.2) to ensure reliable comparison.. In line with the previous findings, all 3 KMT mutant mice exhibit a substantial number of TSS regions (+/− 2kb) with decreased H3K4me3 (adjusted p-value < 0.1 & fold change < −1.2) (**Fig 4A; Supplementary Tables 13-15**). Interestingly, we also detected TSS regions with decreased H3K4me1 in all mutant mice, but only in *Setd1b* cKO mice a substantial number of TSS regions exhibited increased H3K4me1 (**Fig 4B; Supplementary Tables 16-18**). In addition, there was little overlap amongst the TSS regions with decreased H3K4me3 in *Kmt2a*, *Kmt2b* and *Setd1b* cKO mice, providing further evidence that *Setd1b* controls a unique gene expression program in neuronal cells (**Fig 4C**). To test this hypothesis directly, we decided to compare the corresponding gene expression changes in the hippocampal CA region of the 3 different KMT cKO mice. While the ChIP-seq data available for *Kmt2a* and *Kmt2b* cKO mice had been generated from neuronal nuclei, the corresponding gene expression analysis represents RNA-seq data obtained from bulk tissue of the hippocampal CA region (Kerimoglu *et al.*, 2017). To ensure optimal comparison of these RNA-seq data to the gene expression changes in *Setd1b* cKO mice, we also performed RNA-seq from the whole tissue of the hippocampal CA region of *Setd1b* cKO mice and control littermates. We observed 486 genes that were significantly downregulated (adjusted p-value < 0.1 & fold change < −1.2) when comparing control to *Setd1b* cKO mice (**Fig 4D; Supplementary Table 19**). The majority of those genes were also detected via neuronal specific RNA-seq in *Setd1b* cKO mice that we performed before (see above) (**Fig EV9**). While the total number of genes downregulated in the hippocampal CA region of *Kmt2a*, *Kmt2b* and *Setd1b* cKO mice was comparable, there was little overlap amongst them (**Fig 4E; Supplementary Tables 19-21**). Recently, RNA-sequencing data was reported for mice that were heterozygous for *Setd1a*. Although these mutants were heterozygous constitutive knock-out mice and furthermore cortical tissue was analyzed instead of the hippocampus (Mukai *et al*, 2019), it is interesting to note that there was virtually no overlap regarding the genes downregulated in *Setd1a* knock-out mice, when compared to the data obtained from our *Setd1b* cKO mice (**Fig. EV10**). Further support for a distinct role of *Setd1b* in neuronal gene expression was revealed by the finding that the genes downregulated in *Setd1b* cKO mice exhibited a significant enrichment for neuron-specific genes, while this was not the case for genes downregulated in *Kmt2a* cKO or *Kmt2b* cKO mice (**Fig 4F; Fig EV11**). While we observed that genes downregulated in *Kmt2a* or *Kmt2b* cKO mice also display decreased H3K4me3 at the TSS (**Fig 4G**), only the genes downregulated in *Setd1b* cKO mice were characterized by reduced H3K4me3 and also increased H3K4me1 at their TSS (**Fig 4G**). Consequently, the genes that concomitantly exhibited decreased H3K4me3 levels and were downregulated in *Setd1b* cKO mice displayed significantly broader H3K4me3 peaks at the TSS (**Fig 4H**) and were expressed at higher levels under basal conditions when compared to the genes controlled by *Kmt2a* or *Kmt2b* (**Fig 4I; Supplementary Tables 22 and 23**).

**Figure 4.**
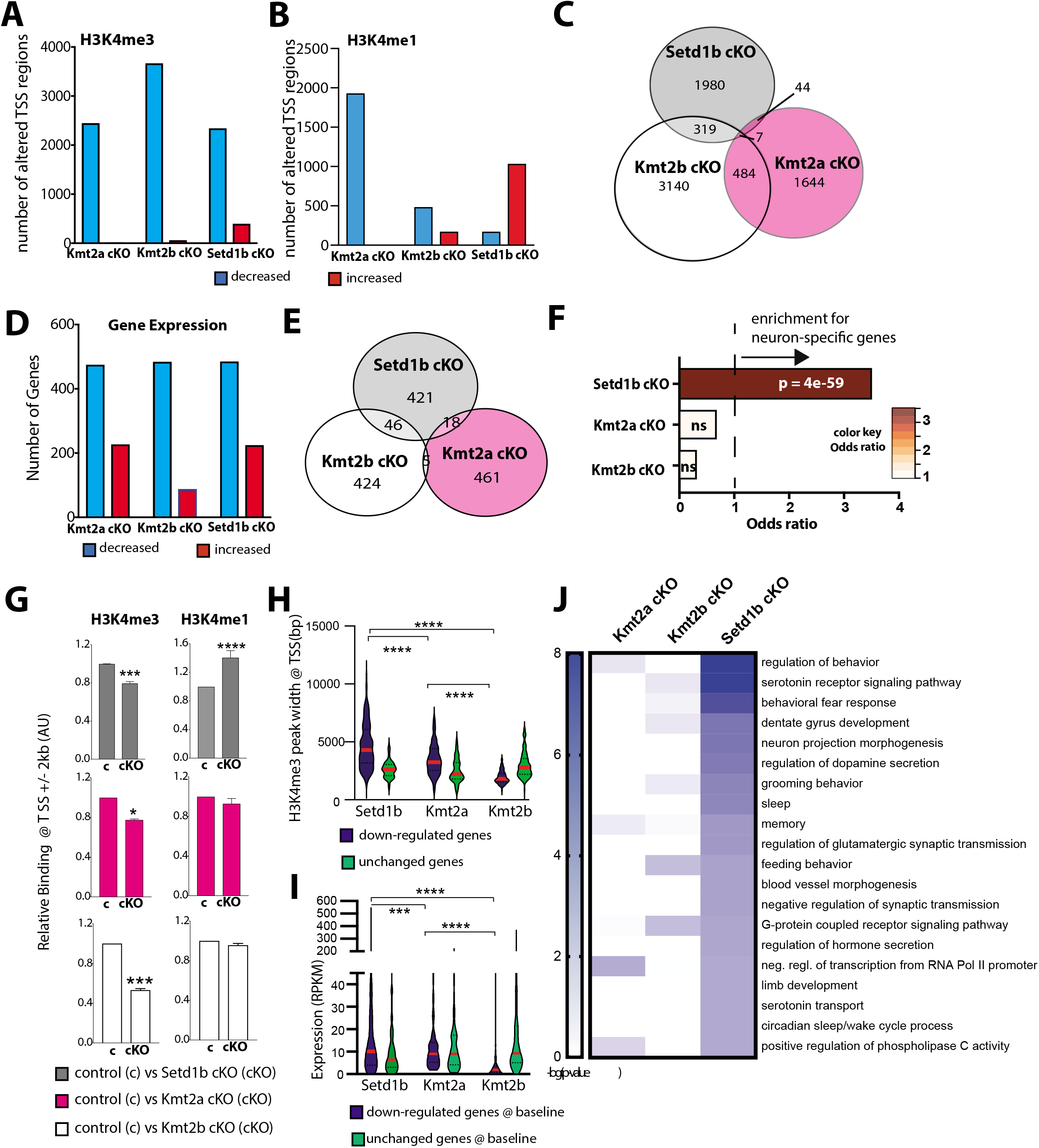
Comparative analysis of the hippocampal epigenome and transcriptome in *Setd1b*, *Kmt2a* and *Kmt2b* cKO mice. **A.** Bar chart showing the number of TSS regions (+/− 2kb) that exhibit significantly altered H3K4me3 (Kmt2a: control, n = 5; cKO, n = 3. Kmt2b: control, n = 6; cKO, n = 5. Setd1b: control, n = 4; cKO, n = 4). **B.** Bar chart showing the number of TSS regions (+/− 2kb) that exhibit significantly altered H3K4me1. **C.** Venn diagram comparing the TSS regions with significantly decreased H3K4me3 in the 3 respective cKO mice. **D.** Bar chart showing the number of differentially expressed genes from bulk RNA-seq in each of the 3 KMT cKO mice. Kmt2a: control, n = 5; cKO, n = 6. Kmt2b: control, n = 8; cKO, n = 11. Setd1b: control, n = 6; cKO, n = 6). **E.** Venn diagram comparing the significantly downregulated genes amongst the 3 respective cKO mice. **F.** Downregulated genes with concomitantly decreased H3K4me3 in each of the 3 KMT cKO mice were tested for the overlap to the 836 neuron-specific genes we had defined for the hippocampal CA region (See Fig EV5). Only for *Setd1b* cKO mice a highly significant odds ratio (Fisher’s exact test) was observed (65 out of 161), while there was no significant association amongst neuron-specific genes and the genes affected in *Kmt2a* cKO (9 out of 112) or *Kmt2b* cKO (20 out of 379) mice (see also Fig EV11 for all downregulated genes in each of the KMT cKOs). **G.** Left panel: Bar graphs showing H3K4me3 binding around the TSS (+/− 2kb) of downregulated genes in either of the 3 KMT cKO mice (Two-way ANOVA: * *p*-value < 0.05, *** *p*-value < 0.001). Right panel depicts H3K4me1 binding at the same TSS regions (Two-way ANOVA: **** *p*-value < 0.0001). Note that only in *Setd1b* cKO mice decreased H3K4me3 is accompanied by significantly increased H3K4me1. **H.** Genes exhibiting decreased H3K4me3 and reduced expression in *Kmt2a*, *Kmt2b* or *Setd1b* cKO mice were analyzed for H3K4me3 peak width around the TSS under basal conditions. Genes affected in *Setd1b* cKO mice displayed significantly broader H3K4me3 peak width when compared to genes affected in *Kmt2a* or *Kmt2b* cKO mice. H3K4me3 peak widths at random sets of unchanged genes of equal size are shown for comparison. Comparisons between genes with decreased H3K4me3 and expression in the KMT cKOs: One-way ANOVA, *p*-value < 0.0001. Post-hoc multiple comparisons, Tukey’s test: *Setd1b* cKO vs *Kmt2a* cKO, **** *p*-value < 0.0001; *Setd1b* cKO vs *Kmt2b* cKO, **** *p*-value < 0.0001; *Kmt2a* cKO vs *Kmt2b* cKO, **** *p*-value < 0.0001. Comparisons between “down” and “unchanged” sets in each KMT cKO: *Setd1b* cKO, t-test *p*-value < 0.0001; *Kmt2a* cKO, Student t-test *p*-value < 0.0001; *Kmt2b* cKO, Student t-test *p*-value < 0.0001. **I.** Violin plots showing average basal expression of downregulated genes with decreased H3K4me3 levels at the TSS in *Kmt2a*, *Kmt2b* or *Setd1b* cKO mice. Genes affected in *Setd1b* KO mice are expressed at significantly higher levels under basal conditions when compared to genes affected in *Kmt2a* cKO or *Kmt2b* cKO mice. Comparisons between genes with decreased H3K4me3 and expression in the KMT cKOs: One-way ANOVA, *p*-value < 0.0001. Post-hoc multiple comparisons, Tukey’s test: *Setd1b* cKO vs *Kmt2a* cKO, *p*-value = 0.1180; *Setd1b* cKO vs *Kmt2b* cKO, **** *p*-value < 0.0001; *Kmt2a* cKO vs *Kmt2b* cKO, **** *p*-value < 0.0001. Comparisons between “down” and “unchanged” sets in each KMT cKO: *Setd1b* cKO, t-test *p*-value < 0.01; *Kmt2a* cKO, t-test *p*-value = 0.3967; *Kmt2b* cKO, t-test *p*-value < 0.0001. **J.** Heat map showing functional pathways of genes affected in *Kmt2a*, *Kmt2b* or *Setd1b* cKO mice. Note that genes affected by loss of Setd1b specifically represent pathways linked to excitatory neuronal function. Error bars indicate SEM.

The fact that under basal conditions genes regulated by *Setd1b* exhibit a wider H3K4me3 distribution at the TSS compared to those regulated by the other two KMTs suggests that *Setd1b* spends more time at the TSS and therefore moves further downstream, which is in line with the suggested mode of action for H3K4 KMTs (Soares *et al.*, 2017). This view is further supported by our observation that the number of genomic regions with decreased H3K4me3 was similar in *Kmt2a*, *Kmt2b* and *Setd1b* cKO mice when we analyzed the first 2 kb downstream of the TSS (**Fig EV12A, B**). However, significantly more regions were affected in *Setd1b* cKO mice when we analyzed the region 5 kb downstream the TSS (**Fig EV12B**). In turn, the genomic regions showing decreased H3K4me3 in *Kmt2a* cKO and *Kmt2b* cKO are concentrated in very close vicinity of TSS when compared to the regions affected in *Setd1b* cKO mice (**Fig EV12C-E**). These data also support the view that *Setd1b* promotes efficient tri-methylation of H3K4 at genes that already underwent mono-methylation by other KMTs.

In line with our data showing that the genes affected in *Kmt2a*, *Kmt2b* and *Setd1b* cKO mice substantially differ, functional GO category analysis revealed that genes affected in the 3 KMT cKO mice also represent different functional categories. When compared to the genes affected in *Kmt2a* or *Kmt2b* cKO mice, genes that were downregulated and concomitantly exhibited decreased H3K4me3 in *Setd1b* cKO mice represent pathways intimately linked to learning and memory and the function of hippocampal excitatory neurons such as “regulation of behavior”, or “memory” (**Fig 4J; Supplementary Table 24**). Further analysis revealed that the genes affected in *Kmt2a* cKO mice also partly relate to neuronal plasticity functions (**Fig EV13A; Supplementary Table 24**) while the genes affected in *Kmt2b* cKO mice represent almost exclusively pathways important for basal cellular function such as metabolic processes (**Fig EV13B; Supplementary Table 24**). We also compared the genes with significantly reduced H3K4me3 at the TSS region (+/− 2 kb) among the three KMT cKOs and a study that analyzed H3K4me3 in bulk hippocampus tissue of a mouse model for Alzheimer’s disease (Gjoneska *et al.*, 2015). Interestingly, the overlap was significantly higher for *Setd1b* cKO mic (**Fig EV14**).

In summay, our observations so far support the view that the *Setd1b*, *Kmt2a* and *Kmt2b* serve distinct functions in excitatory hippocampal neurons oft the postnatal brain and that among them *Setd1b* might be the most relevant for the regulation of learning processes. In line with this, comparison of hippocampus-dependent learning in *Setd1b*, *Kmt2a* and *Kmt2b* cKO mice in the Morris water maze task, suggest that learning impairment is more pronounced in *Setd1b* cKO mice (**Fig EV15A**). To further elucidate the distinct roles of *Setd1b*, *Kmt2a* and *Kmt2b* we hypothesized that they might be expressed in different neuronal subtypes. Therefore, we isolated the hippocampus from 3-month-old wild type mice, sorted the nuclei and performed single nuclear (snuc) RNA-seq (**Fig EV15B**). We were able to detect all major cell types of the hippocampus and found *Kmt2a*, *Kmt2b* and *Setd1b* as well as the other three H3K4 KMTs to be expressed in all of these cell types (**Fig EV15C**). We did not observe any particular neuronal cell type with especially high *Setd1b* expression (**Fig EV15D**). Interestingly, the snucRNA-seq data revealed higher *Kmt2a* expression when compared to *Setd1b* and *Kmt2b* across cell types including excitatory neurons of the CA region, an effect that was not due to the total amount of reads detected per cell (**Fig EV15C, D**). These data were confirmed via RNAscope. Thus, when we analyzed the pyramidal neurons of the hippocampal CA region, we observed significantly more transcripts/cell for *Kmt2a* when compared to *Kmt2b* or *Setd1b* (**Fig EV15E**). In summary, these data may suggest that highly expressed enzymes such as *Kmt2a* are essential to ensure neuronal H3K4 methylation sufficient for basal and neuron-specific cellular processes and that the additional expression of enzymes such as *Setd1b* is essential to further promote H3K4me3 at genes particularly important for neuronal functions like synaptic plasticity.

## DISCUSSION

We show that postnatal loss of *Setd1b* from excitatory forebrain neurons impairs learning in mice when measured in the hippocampus-dependent water maze paradigm. These data are in line with previous findings showing that hippocampal H3K4me3 increases in response to memory training in rodents (Gupta *et al.*, 2010), while its levels are reduced in the hippocampus of a mouse for AD-like neurodegeneration (Gjoneska *et al.*, 2015) and in postmortem human brain samples of patients suffering from cognitive diseases (Shulha *et al*, 2012). Our data furthermore support previous genetic studies linking mutations in *Setd1b* to intellectual disability (Labonne *et al.*, 2016) (Hiraide *et al.*, 2018) (Roston *et al.*, 2020). Although spatial reference memory measured in the water maze paradigm critically depends on hippocampal function, it is important to note that *Setd1b* deletion is not specific to the hippocampus and that we cannot exclude that other brain regions and subtle changes in postnatal development may also contribute to the observed phenotype. Impaired hippocampus-dependent memory has also been observed in mice that lack *Kmt2a* (Gupta *et al.*, 2010) (Kerimoglu *et al.*, 2017) or *Kmt2b* (Kerimoglu *et al.*, 2013) from excitatory neurons of the postnatal forebrain. Our comparative analysis suggests, however, that the effect is most severe in *Setd1b* cKO mice. Interestingly, mice heterozygous for *Setd1a*, the close homologue to *Setd1b* that is genetically linked to schizophrenia, show no impairment in the water maze task but rather exhibit impaired working memory and schizophrenia-like phenotypes (Mukai *et al.*, 2019). Distinct roles for *Setd1b* and *Setd1a* are also reported for other cellular system (Bledau *et al*, 2014) (Arndt *et al*, 2018; Brici *et al*, 2017; Kranz & Anastassiadis, 2020; Schmidt *et al*, 2018). These data suggest that the different H3K4 KMTs, and at least *Kmt2a*, *Kmt2b*, *Setd1a* and *Setd1b*, likely serve distinct functions in the postnatal brain. It will therefore be important to eventually generate corresponding and comparable data for all six H3K4 KMTs.

The molecular characterization of *Setd1b* cKO further confirms this view. Here, we focused on the analysis of the hippocampus, since this brain region is critical for spatial reference memory which was severely impaired in *Setd1b* cKO mice. It will nevertheless be interesting to study other brain regions such as the prefrontal cortex in future experiments. In line with the role of *Setd1b* in regulating H3K4me3, we observed a substantial decrease of neuronal H3K4me3 levels and the vast majority of these changes were observed close to TSS regions. Many genes with decreased H3K4me3 also exhibited reduced H3K9ac, which is in line with previous data showing that H3K4me3 appears to be a pre-requisite for H3K9ac, most likely since H3K4 KMTs interact with histone acetyltransferases (Kerimoglu *et al.*, 2013) (Mishra *et al*, 2014) (Wang *et al*, 2009). For example, SETD1B and the histone-acetyltransferase KAT2A were shown to interact with WDR5, a central component of H3K4 KMT complexes (Lin *et al*, 2016) (Ma *et al*, 2018), which is interesting since loss of *Kat2a* from excitatory forebrain neurons also leads to severe impairment of spatial reference memory (Stilling *et al.*, 2014).

Our finding that H3K4me1 levels were increased at a substantial number of TSS regions that exhibited decreased H3K4me3 in *Setd1b* cKO mice suggests that H3K4 mono-methylation at the TSS regions of the affected genes is likely mediated by other KMTs in addition to *Setd1b*. This is in agreement with increasing evidence suggesting that the different H3K4 KMTs exhibit some specificity towards mono- di- or tri-methylation. However, specificity likely depends on the cellular context and care has to be taken when interpreting the different *in vitro* and *in vivo* data. Nevertheless, our data are in line with studies showing that SETD1B preferentially mediates H3K4-trimethylation when compared to the other KMTs such as KMT2A and KMT2B that are believed to mediate mono- and di-methylation with comparatively low H3K4 tri-methylation activity (Wu *et al*, 2008) (Lee & Skalnik, 2008) (Shinsky *et al*, 2015) (Bochyńska *et al.*, 2018; Rao RC, 2015) (Kranz & Anastassiadis, 2020).

In agreement with these findings, we show that neuronal H3K4me1 is exclusively reduced in *Kmt2a* and *Kmt2b* cKO. Since H3K4me3 is similarly reduced in these cKO mice, our data suggest that neuronal KMT2A and KMT2B can also catalyze H3K4 tri-methylation. In such a scenario, SETD1B might act in concert with other KMTs to further facilitate H3K4me3 at specific genes. This might also explain our finding that genes affected in *Setd1b* cKO mice exhibit the widest H3K4me3 distribution at their TSS and the highest RNA expression levels under basal conditions. A broad H3K4me3 peak around TSS is indicative of higher transcriptional frequency and RNA expression levels and has thus been linked to the expression of genes that are of particular importance for a given cell type (Benayoun *et al.*, 2015) (Park S, 2020). This is in line with our observation that genes decreased in hippocampal neurons in *Setd1b* cKO mice are characterized by a broad H3K4me3 peak and represent functional pathways critically involved in synaptic plasticity and learning & memory. In agreement with this a recent study showed that memory training specifically activates hippocampal genes with broad H3K4me3 peaks at the TSS (Collins *et al*, 2019). It is therefore interesting to note that the genes affected in *Setd1b* cKO mice not only showed a broader H3K4me3 peak and higher expression when compared to the genes affected in *Kmt2a* or *Kmt2b* cKO mice but that also memory performance in the water maze training was more severely affected in *Setd1b* cKO mice when compared to mice lacking *Kmt2a* or *Kmt2b*. These data are also in line with a recent study in mouse embryonic stem cells in which SETD1B was associated with the regulation of highly expressed genes that exhibit a broad H3K4me3 peak, while KMT2B was linked to the expression of genes with narrow H3K4me3 peaks (Sze *et al*, 2020). Interestingly, that study suggested a functional redundancy of *Setd1b* and *Setd1a*. However, our data suggest that this is not the case in post-mitotic neurons. Moreover, mutations in either *Setd1a* or *Setd1b* lead to distinct neuropsychiatric diseases and unlike *Setd1b* cKO mice, *Setd1a* heterozygous mutant mice do not exhibit impairment of long-term memory consolidation (Mukai *et al.*, 2019).

These findings may suggest that among the three H3K4 KMTs studied here, *Kmt2a* and *Kmt2b* are essential to ensure the sufficient expression of genes important for basal cellular processes and in the case of *Kmt2a* also specific neuronal functions. The additional presence of *Setd1b* may further facilitate H3K4me3, thereby enabling the preeminent expression of genes specific for hippocampal excitatory neuronal function.

At present we cannot conclusively answer the question how *Setd1b* affects the expression of a specific set of hippocampal genes. Our snucRNA-seq analysis showed that *Setd1b* is expressed at similar levels in all types of excitatory neurons of the hippocampal CA region suggesting that its specific function of facilitating learning behavior is not driven by its expression in a specific neuronal subtype. The fact that loss of either *Kmt2a*, *Kmt2b* or *Setd1b* leads to distinct changes in neuronal gene-expression might also be due to the specific subunit compositions of the KMT complexes. Previous data suggest that H3K4 KMTs associate with different co-activators (Yokoyama *et al*, 2004) (Scacheri *et al*, 2006) (Lee *et al*, 2006) (Dreijerink *et al*, 2006) (Shilatifard, 2012) (Bochyńska *et al.*, 2018) (Kranz & Anastassiadis, 2020). While all H3K4 KMTs are believed to interact with a number of core subunits (Kranz & Anastassiadis, 2020), data from other cell types than neurons show that KMT2A and KMT2B can bind the transcriptional regulator multiple endocrine neoplasia type 1 (*Menin*) (Hughes *et al*, 2004). In contrast, SETD1B was shown to interact with CXXC finger protein 1 (CFP1) and WDR82 which binds to RNA-polymerase II. This is in line with data suggesting that SETD1A and SETD1B are the major KMTs found at TSS regions (Bi *et al*, 2011; Deng *et al*, 2013).

Future studies will need to further explore this possibility. It is however, interesting to note that the genes with decreased H3K4me3 in the hippocampus of *Kmt2a* or *Kmt2b* cKO mice were enriched for binding sites of ETS and E2F transcription factors, respectively (Kerimoglu *et al.*, 2017).

In case of *Setd1b* cKO mice we observed an enrichment for REST, a key transcription factor inhibiting the expression of neuron-specific genes (Hwang & Zukin, 2018). This observation may help to understand why loss of *Setd1b* mainly affected the expression of neuron-specific genes important for memory function. While future research should aim to further elucidate the specific role of SETD1B, it is interesting to note that the H3K4 Demethylase KMD5C was found to regulate REST target genes (Tahiliani *et al*, 2007).

Taking into account that decreased neuronal H3K4me3 levels have been observed in cognitive and neurodegenerative diseases, therapeutic strategies that reinstate specifically the expression of neuronal plasticity genes controlled by SETD1B might be particularly helpful. We suggest that the epigenetic drugs currently tested in pre-clinical and clinical settings for cognitive diseases should also be analyzed for their potential to reinstate the H3K4me3 peak width at the TSS of genes linked to neuronal function and learning behavior.

The aim of this study was to analyze the role of *Setd1b* on learning behavior when deleted postnatally and to explore its impact on epigenetic gene expression and compare these data to findings obtained from postnatal deletion of *Kmt2a* and *Kmt2b*. While we observe that *Setd1b* is required for memory function and controls the expression of genes linked to neuronal plasticity, it will be important to study the functional and structural consequences in the corresponding neuronal networks. At present such data is rare. A recent study employed heterozygous *Kmt2a* constitutive knockout mice and found decreased dendritic spine density in the ventral hippocampal CA1 region (Vallianatos *et al*, 2020). To compare the role of the different H3K4 KMTs in synaptic morphology and network activity during development and in the postnatal brain is thus an important task for future research.

In summary, we identify H3K4 methyltransferase *Setd1b* as a crucial factor regulating genes important for hippocampal excitatory neuronal function, which are involved in synaptic plasticity and learning.

## MATERIALS AND METHODS

### Animals

All animals used in this study were of the same genetic background (C57BL/6J) mice and of 3-6 months of age. The experimental groups were age and sex matched. Mice were kept in standard home cages with food and water provided *ad libitum*. All experiments were performed according to the animal protection law of the state of Lower Saxony. The CRE mice were first described by Minichiello et al (Minichiello *et al.*, 1999) and were used in previous studies related to *Kmt2a* and *Kmt2b* (Kerimoglu *et al.*, 2013) (Kerimoglu *et al.*, 2017). Deletion of the target gene is restricted to the forebrain excitatory neurons and is initiated upon postnatal day 21.

### Behavior experiments

#### Water maze

The behavioral experiments were performed as described previously (Kerimoglu *et al.*, 2017). For in depth feature analysis from water maze data, a modified version of MUST-C algorithm was used (Illouz *et al.*, 2016) (Islam *et al.*, 2021). In brief, different strategies were defined based on trajectories mice employed in each trial. Cumulative score for hippocampal dependent strategy score was calculated as follow:

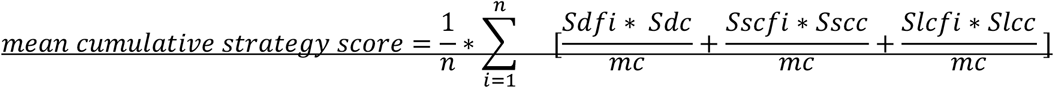

n=total trial number

Sdf_i_ = Frequency of direct strategy in i^th^ trial

Sdc = given strategy score for direct search; 10

mc = total number of mice

Sscf_i=_ Frequency of short chaining in i^th^ trial

Sscc = given strategy score for short chaining search; 9.5

Slcf_i=_ Frequency of short chaining in i^th^ trial

Slcc = given strategy score for long chaining search; 7

#### Y-maze & Open field

The open field tests were performed as described previously (Islam *et al.*, 2021). Working memory was accessed with a Y-shaped plastic runway (10 x 40 cm arms, walls 40 cm high). The mouse was placed into the Y-maze in the triangle-shaped central platform and left to freely explore the maze for 5 min. % of spontaneous alterations (choice of “novel” arm: when animal goes into different arm then before) was recorded and scored via a videosystem (TSE).

### Tissue isolation and processing

ChIP-Seq and bulk RNA-Seq experiments from NeuN (+) nuclei were performed from the hippocampal cornu ammonis (CA) region, which consists of the hippocampus without the dentate gyrus and was isolated as described before (Kerimoglu *et al.*, 2013) (Kerimoglu *et al.*, 2017). For single nuclear RNA-sequencing whole hippocampal tissue was isolated. The tissues were dissected and flash frozen in liquid nitrogen and stored at −80°C until further processing.

### Cell-type specific nuclear RNA isolation and sequencing

Frozen CA tissues from the left and right hemisphere of two mice were pooled together and processed on ice to maintain high RNA integrity. Tissue was homogenized using a plastic pestle in a 1.5mL Eppendorf tube containing 500 uL EZ prep lysis buffer (Sigma, NUC101-1KT) with 30 strokes. The homogenate was transferred into 2 mL microfuge tubes, lysis buffer was added up to 2 mL and incubated on ice for 7 minutes. After centrifuging for 5 minutes at 500g, the supernatant was removed and the nuclear pellet was resuspended into 2 mL lysis buffer and incubated again on ice (7 minutes). After centrifuging for 5 minutes at 500g, the supernatant was removed and the nuclei pellet was resuspended into 500ul nuclei storage buffer (NSB: 1x PBS; Invitrogen, 0.5% RNase free BSA;Serva, 1:200 RNaseIN plus inhibitor; Promega, 1x EDTA-free protease inhibitor; Roche) and filtered through 40 μm filter (BD falcon) with additional 100 μL NSB to collect residual nuclei from the filter. Nuclei were stained with anti-NeuN-Alexa488 conjugated antibody (1:1000) for 45 minutes and washed once with NSB. Stained nuclei were then FACS-sorted with FACSaria III using 85 μm nozzle. Nuclei were gated by their size, excluding doublets and neuronal nuclei were separated from non-neuronal nuclei by their NeuN-Alexa488 fluorescence signal. Sorted nuclei were collected into a 15 mL falcon tube precoated with NSB, spun down and RNA was isolated using Trizol LS. After addition of chloroform according to the Trizol LS protocol, aqueous phase was collected and RNA was isolated by using Zymo RNA clean & concentrator-5 kit with DNAse treatment. Resulting RNA concentration were measured in Qubit and RNA-seq was performed using 100ng of neuronal RNA with illumina TruSeq RNA Library Prep Kit. Since glial nuclei are smaller and contains very little amount of RNA, neuronal nuclear RNA was scaled down and 1ng from both neuronal and glial nuclear RNA was used to make RNA-seq libraries using Takara SMART-Seq v4 Ultra Low Input RNA Kit. Libraries were sequenced using single-end 75 bp in Nextseq 550 or single-end 50 bp in HiSeq 2000, respectively.

### Cell-type specific chromatin isolation and ChIP sequencing

Frozen tissues were homogenized, formaldehyde (1%) fixed for 10 minutes and quenched with 125mM glycine for 5 minutes. Debris was removed by sucrose gradient centrifugation. The resulting nuclear pellet was stained with anti-NeuN-Alexa488 conjugated antibody (1:1000) for 25 minutes and washed 3 times with PBS. Stained nuclei were then FACS sorted with FACSaria III using 85 μm nozzle. Nuclei were gated similarly as described previously (Halder *et al*, 2016). Sorted nuclei were collected into a 15mL falcon tube and transferred into 1.5mL tubes. The nuclear pellet was flash frozen in liquid nitrogen and saved at −80°C for further processing. For chromatin shearing, the pellet was resuspended into 100uL RIPA buffer (containing 1% SDS) and sonicated for 25 cycles in Diagenode bioruptor plus with high power and 30 cycles on/ 30 cycles off. Chromatin shearing was checked by taking a small aliquot and decrosslinking the DNA by 30 minutes RNAse and 2 hours of proteinase K treatment. DNA was isolated using SureClean Plus protocol. Sheared chromatin size was determined using Bioanalyzer 2100(DNA high sensitivity kit) and the concentration was measured using Qubit 2.0 fluorometer (DNA high sensitivity kit). 0.3μg chromatin was used along with 1 μg of antibody to do ChIP for H3K4me3 (Abcam ab8580), H3K4me1 (Abcam ab8895), H3K27ac (Abcam ab4729) and H3K9ac (Millipore 07-352). ChIP was performed as previously described (Halder *et al.*, 2016). The resulting ChIP DNA was subjected to library preparation using NEBNext Ultra II DNA library preparation kit and sequenced for single end 50 bp at illumina HiSeq 2000.

### ChIP-Seq Analysis

#### Pre-Processing, Profile Plots, Peak Calling, Differential Binding Analysis, Annotation, Motif Enrichment Analysis and Visualization

Base calling and fastq conversion were performed using Illumina pipeline. Quality control was performed using fastqc (https://www.bioinformatics.babraham.ac.uk/projects/fastqc). Reads were mapped to mm10 mouse reference genome with STAR aligner v2.3.0.w. PCR duplicates were removed by *rmdup -s* function of samtools (Li *et al*, 2009). BAM files with unique reads belonging to the same group were merged into a single BAM file with the *merge* function of samtools. Profile plots were created from these merged BAM files with NGSPlot (Shen *et al*, 2014).

Peak calling was performed using MACS2 (Feng *et al*, 2012) against the input corresponding to the particular group (i.e., control or cKO) using q < 0.1. Consensus peaksets were generated for each histone modification individually using the Diffbind package of Bioconductor (Ross-Innes *et al*, 2012) with the command *dba.count* and the parameter *minOverlap=1*. Then, these consensus peaksets were intersected with each other using the *intersect* function of bedtools with default parameters, providing one common peakset representing all four histone modifications. The differential binding analysis for each histone mark between control and *Setd1b* cKO was then performed using Diffbind (Ross-Innes *et al.*, 2012) with this common peakset as input.

For the comparison of H3K4me3 changes in *Kmt2a* cKO, *Kmt2b* cKO and *Setd1b* cKO common peaksets for each individual histone mark from three separate ChIP-Seq experiments were extracted in the same way as above. In this case, first, consensus peaksets for a histone mark from each individual ChIP-Seq experiment (i.e., “Control vs *Kmt2a* cKO”, “Control vs *Kmt2b* cKO” and “Control vs *Setd1b* cKO”) were determined using Diffbind as above. Then, these consensus peaksets were intersected with each other as above, and as a result we obtained a set of common regions detected in all three independent experiments that possess H3K4me3 mark. For the purpose of comparing the effects of the three KMT knockdowns on H3K4me3, the differential binding analysis for each individual ChIP-Seq experiment was performed using this common peakset as input.

We also wanted to compare the changes in H3K4me1 in these three KMT cKO mice. But additionally, based on our prior observations from *Setd1b* cKO mice, we wanted to also investigate the interplay between H3K4me3 and H3K4me1, and whether it differs between the three KMT cKO mice. We therefore first came up with the common peakset for H3K4me1 from all three ChIP-Seq experiments as described above. Then, we intersected this common H3K4me1 peakset with the common H3K4me3 peakset and came up with a peakset representing regions containing both H3K4me3 and H3K4me1 marks in all three separate experiments (“me3_me1_3_kmt_ckos”). The differential binding analyses and comparisons relating to H3K4me1 in the three KMT cKOs were performed using “me3_me1_3_kmt_ckos” as input. Diffbind package was used for differential binding analysis with in-built DESEQ2 option for differential analysis (Ross-Innes *et al.*, 2012). For the initial analysis of *Setd1b* cKO alone we used the cut-off “adjusted p-value < 0.05 & |fold change| > 1.5” to determine significance. For comparing differential binding in *Kmt2a* cKO, *Kmt2b* cKO and *Setd1b* cKO mice we used a more lenient cut-off – “adjusted p-value < 0.1 & |fold change| > 1.2” in order to avoid missing out on overlaps and/or common trends that might be biologically relevant. The annotation of the genomic regions and transcription factor motif enrichment analysis were performed with HOMER (Heinz *et al*, 2010). ChIPSeeker package of Bioconductor (Yu *et al*, 2015) was used to calculate the proportion of the types of genomic regions in a given set and to create corresponding pie charts. Graph Pad Prism was used to generate violin plots. Outliers were removed using the Rout methods (Q = 1%) in GraphPad Prism. Integrated Genome Browser (IGB) was used to make visualizations representing the distribution of histone marks and RNA expression over selected genes (Nicol *et al*, 2009).

### Obtaining H3K4me3 Width at Promoters

#### 1. For Promoters with Decreased H3K4me3 and Different H3K4me1 Trends at TSS Proximal Regions in Setd1b cKO

First, we performed differential binding analysis in “Control vs *Setd1b* cKO” for H3K4me3 and H3K4me1 using the “me3/me1/k9ac/k27ac common peakset”. Then we extracted the TSS proximal regions (+/− 2000 bp from TSS) with significantly decreased H3K4me3 in *Setd1b* cKO (adjusted p-value < 0.05 & fold change < −1.5). These TSS proximal regions were then separated into three groups depending on the concomitant change of H3K4me1; (i) increased H3K4me1 (fold change > 1.5), (ii) decreased H3K4me1 (fold change < −1.5) and (iii) not changed H3K4me1 (the rest). Then each of these TSS proximal regions was intersected to the consensus H3K4me3 peakset with bedtools using *intersect* function with the option “*-u*”. The latter option ensures that the full original feature in the first input (in this case always the H3K4me3 peakset), *not just the overlapping portion*, is reported once the overlap is found. In this way the full extent of H3K4me3 distribution around the target genomic region can be appreciated. Finally, the width was calculated by subtracting chromosome start coordinates from chromosome end coordinates and adding 1 (chromosome end – chromosome start + 1).

#### 2. For Promoters with Decreased H3K4me3 at TSS Proximal Regions in Different KMT cKOs

For the purpose of comparison between the three KMT cKO mice we chose a more lenient cut-off (adjusted p-value < 0.1 & fold change < −1.2) in order to avoid as much as possible excluding genomic regions showing the same trend but not making it past the threshold in some of the cases. The TSS proximal regions (+/− 2000 bp) with significantly decreased H3K4me3 in each KMT cKO were overlapped to the consensus H3K4me3 peakset using *intersect –u*. Again, in this way we were able to capture the full extent of H3K4me3 around the selected target regions. The width at the resulting region was then calculated by chromosome end – chromosome start + 1. A random list of the same size was also generated from the TSS proximal regions with unchanged H3K4me3 for each cKO using the same procedure. The criteria for unchanged H3K4me3 were adjusted p-value > 0.5 & |log2(fold change)| < 0.1.

#### 3. For Downregulated Genes

Again, to obtain a more comprehensive view of downregulation of gene expression and its concordance with peak width we chose a more lenient cut-off; adjusted p-value < 0.1 & fold change < −1.2.

From the consensus H3K4me3 peakset the promoter regions were selected and annotated for their gene names using HOMER. To capture the whole extent of H3K4me3 along the promoters as much as possible, we used the default standard for promoter identification implemented by ChIPSeeker (+/− 5000 bp; see above). From the resulting annotated file the promoter regions belonging to significantly downregulated genes were extracted and the widths were calculated as above. In each case, a random list of the same size from genes with unchanged expression was also generated using the same procedure. The criteria for unchanged expression were adjusted p-value > 0.5 & |log2(fold change)| < 0.1.

#### 4. For the Genes with Both Decreased TSS Proximal H3K4me3 and Downregulated Expression in Different KMT cKOs

The bed files generated in Section 2 by the overlap of the consensus H3K4me3 peakset with TSS proximal regions having significantly decreased H3K4me3 were annotated for gene names using HOMER. From the resulting annotated file the genomic coordinates belonging to the downregulated genes were extracted and the widths were calculated. Again, in each case a random list of the same size containing genes with unchanged expression and H3K4me3 was selected using the same procedure and the same criteria as above.

### RNA-Seq Analysis

Base calling, fastq conversion, quality control, mapping of reads were performed as described before (Kerimoglu *et al.*, 2017). Bulk RNA-Seq was mapped with STAR on the whole genome and reads were counted with featureCounts thus considering spliced as well as unspliced transcripts. Differential expression was analyzed with DESeq2 package of Bioconductor (Love *et al*, 2014). RPKM values were calculated using edgeR package of Bioconductor. We used the cut-off “adjusted p-value < 0.1 & |fold change| > 1.2” to determine significance.

### Functional GO Enrichment Analysis

Enrichment for functional GO categories was performed with the topGO package of Bioconductor (https://bioconductor.org/packages/release/bioc/html/topGO.html), using the weighted analysis option as described in the manual with default settings. As a result, a weighted p-value for each GO category was calculated (described in the manual). In each separate analysis only the GO categories (biological processes) with a weighted *p*-value lower than 0.005 were selected and shown in the figures. All genes in the genome were used as the reference set (i.e., gene universe). The whole list of the GO categories, including the ones highlighted in the figures, can be found in supplemental figures 7, 12 and 24.

### Single-nucleus RNA-Seq

Unfixed NeuN+ neuronal nuclei were isolated as mentioned above (section: Cell-type specific nuclear RNA isolation and sequencing). Sorted neuronal nuclei were counted in a Neubauer chamber with 10% trypan blue (in PBS) and nuclei concentration were adjusted to 1000 nuclei/μL. The nuclei were further diluted to capture and barcode either 2000 or 4000 nuclei according to Chromium single cell 3ʹ reagent kit v3 (10X genomics). Single nuclei barcoding, GEM formation, reverse transcription, cDNA synthesis and library preparation were performed according to 10X genomics guidelines. Finally, the library was sequenced in Illumina NextSeq 550 according to manufacturer’s protocol. Gene counts were obtained by aligning reads to the mm10 genome (GRCm38.p4) (NCBI:GCA_000001635.6) using CellRanger software (v.4.0.0) (10XGenomics). The CellRanger count pipeline was used to generate a gene-count matrix by mapping reads to the pre-mRNA as reference to account for unspliced nuclear transcripts. The CellRanger aggr pipeline was used to generate an aggregated normalized gene-count matrix from seven datasets. The final count matrix contained 15661 cells with a mean of 56.150 total read counts over protein-coding genes.

The SCANPY package was used for pre-filtering, normalization and clustering (Wolf *et al*, 2018). Initially, nuclei that reflected low-quality cells (either too many or too few reads, nuclei isolated almost exclusively, nuclei expressing less than 10% of house-keeping genes (Eisenberg & Levanon, 2013) were excluded remaining in 15656 nuclei. Next, counts were scaled by the total library size multiplied by 10.000, and transformed to log space. A total of 3771 highly variable genes were identified based on dispersion and mean, the technical influence of the total number of counts was regressed out, and the values were rescaled. Principal component analysis (PCA) was performed on the variable genes, and UMAP was run on the top 50 principal components (PCs) (Becht *et al*, 2018). The top 50 PCs were used to build a k-nearest-neighbours cell–cell graph with k= 100 neighbours. Subsequently, spectral decomposition over the graph was performed with 50 components, and the Leiden clustering algorithm was applied to identify nuclei clusters. We confirmed that the number of PCs captures almost all the variance of the data. For each cluster, we assigned a cell-type label using manual evaluation of gene expression for sets of known marker genes. Violin plots for marker genes were created using the stacked_violin function as implemented in SCANPY.

### RNAscope

For the detection of *Setd1b*, *Kmt2a* and *Kmt2b*, the RNAscope® Fluorescent Multiplex Assay (ACD Biosystems) was used according to the manufacturer’s instructions for fresh-frozen tissue. Briefly, mice (n = 2) were sacrificed by cervical dislocation and the brains were removed quickly, flash-frozen using liquid nitrogen and embedded in OCT. Then, 20 μm coronal sections were cut and stored at −80°C until further use. For the pre-treatment, the sections were fixed with cold 10% neutral-buffered formalin at 4°C for 15 min and afterwards dehydrated using 50%, 70% and 100% ethanol. Then, the samples were incubated with Protease IV for 30 min at RT. The probes were hybridized to the tissue for two hours at 40°C using the HybEZ™ Humidifying System (ACD Biosystems). Probes were applied to detect *Setd1b*, *Kmt2a* or *Kmt2b*. Positive and negative control probes were also used to control for signal sensitivity and specificity. The probes were then amplified following the manufacturer’s protocol. *Kmt2a* and *Kmt2b* were labeled with Atto647 and *Setd1b* with Atto550. Images were obtained on a Leica DMi8 microscope using a 40x air objective and the software CellProfiler (Carpenter *et al*, 2006) was used for downstream analysis.

## Supporting information

Expanded View Figure 1

Expanded View Figure 2

Expanded View Figure 3

Expanded View Figure 4

Expanded View Figure 5

Expanded View Figure 6

Expanded View Figure 7

Expanded View Figure 8

Expanded View Figure 9

Expanded View Figure 10

Expanded View Figure 11

Expanded View Figure 12

Expanded View Figure 13

Expanded View Figure 14

Expanded View Figure 15

## Data availability

GEO accession GSE180326 (token available via editor)

## Acknowledgments

This work was supported by the following third party funds to AF: The ERC consolidator grant DEPICODE (648898), the BMBF projects ENERGI (01GQ1421A) and Intergrament (01ZX1314D), and funds from the German Center for Neurodegenerative Diseases FS was supported by the DFG grant SA1005/2-1and funds from the DZNE. AM is a student of the International Max Planck Research School (IMPRS) for Genome Science. SS & RI are students of the IMPRS for Neuroscience.

## Author contributions

AM initiated the project as part of her PhD thesis, performed behavior experiments, performed and analyzed immunohistochemistry and analyzed mutant mice; SS generated cell-type specific ChIP/RNA-seq and single nucleus RNA-seq data and contributed to the analysis of cell type specific RNA-seq, CK analyzed and interpreted ChIP-seq and RNA-seq data and supervised all bioinformatic data analysis, DMF analyzed single nucleus RNA-seq data, LK, XX & EMZ helped with the generation of single nucleus RNA-seq; SS, JPD & GE performed and supervised RNAscope experiments, AK & FAS provided material and analyzed data, AM, SS, CK and AF designed experiments and generated figures. CK and AF wrote the paper.

## Expanded View Figure Legends

**Expanded view Fig 1: Behavioral analysis of mice expressing CamKII-driven Cre recombinase. A.** Transgenic mice expressing Cre under control of the CamKII promoter were subjected to behavior testing (n=8, Cre+) comparing them to wild type mice from the same breeding colony that did not express Cre (n=8; Cre−). No difference was observed in body weight. **B-C.** The distance traveled in the open field test (B) and the time spent in the center of the arena (C) were similar amongst groups. **D.** No difference in the swimming speed was observed amongst groups when subjected to the water maze test. **E.** Escape latency during water maze training was similar in Cre- and Cre+ mice. **F-G.** During the probe test performed after 10 training days, Cre- and Cre+ mice showed similar performance when time spent in the target quadrant (F) and the number of platform crossings (G) were analyzed. Error bars indicate SEM.

**Expanded view Fig. 2. Analysis of memory recall cannot be interpreted in Setd1b cKO mice. A.** 24 h after the completion of the 10 day-training procedure in the water maze paradigm (See Fig 1G) mice were subjected to a probe test. The performance in the probe test, as measured by the time spent in the target quadrant (TQ), is impaired in *Setd1b* cKO mice (Control: n = 15, cKO: n = 15. **** Student t-test < 0.0001). These results have to be interpreted with care, however, since faulty interpretation about memory recall is possible in this experiment since the *Setd1b* cKO mice not even being able to form the memory during the training (See Fig 1G, H). It is moreover important to note that *Setd1b* cKO mice spent even less time in tQ than would be predicted by chance alone. To address the issue we analyzed the swim speed during the entire experiment **(B)** and noticed that the swim speed was similar in control and *Setd1b* cKO mice across the first 5 days of training. There was however a trend for reduced swim speed in *Setd1b* cKO mice from training day 6 to 10. Although we did not detect significant differences in the swim speed when we analyzed the average values across all training days, a more detailed analysis revealed that swim speed was significantly impaired in Setd1b cKO mice at training day 10 (*Student t-test = 0.016) and also during the probe test (**Student t-test = 0.001). These data may help to explain why the *Setd1b* cKO mice appear to avoid the target quadrant during the one trial 60 sec probe test session. The training session consists of a sequence of 4 x 60 sec sessions and mice are always placed into the pool from 4 different locations. During the probe test, that consists of a one trial 60 sec session, animals are always placed in the quadrant opposite to the target quadrant, i.e., the quadrant that is the farthest aways from the target quadrant. While the reason for the reduced swim speed after 9 days of training is unclear, at present these data do not affect the conclusion that *Setd1b* cKO mice exhibit impaired learning ability. Error bars indicate SEM.

**Expanded view Fig. 3: A.** Principal component analysis (PCA) plot from H3K4me3 ChIP-Seq experiment. **B.** PCA plot from H3K4me1 ChIP-Seq experiment. **C.** PCA plot from H3K9ac ChIP-Seq experiment. **D.** PCA plot from H3K27ac ChIP-Seq experiment. **E.** PCA plot from RNA-Seq experiment performed from FACS-sorted neuronal (NeuN+) nuclei. **F.** PCA plot from RNA-Seq experiment performed from bulk CA tissue.

**Expanded view Fig. 4: Decreased H3K9ac and H3K27ac in *Setd1b* cKO mice. A.** Left panel: Heat map showing genomic regions with differentially bound H3K9ac sites in the close vicinity of TSS (+/− 2kb) in neuronal nuclei from *Setd1b* cKO mice and the overall genomic locations of altered H3K9ac levels (Control: n = 4; *Setd1b* cKO: n = 3). Right panel shows the same analysis for H3K27ac (Control: n = 3; *Setd1b* cKO: n = 4). **B.** Bar chart showing the number of genes with significantly decreased and increased H3K9ac and H3K27ac marks at the TSS region (+/− 2kb). Data for H3K4me3 and H3K4me1 are shown for comparison. As expected, the most affected histone mark is H3K4me3. **C.** Venn diagram showing that most of the sites exhibiting decreased H3K9ac at the TSS also exhibit reduced H3K4me3, while this is not the case for H3K27ac. TSS, transcription start site.

**Expanded view Fig. 5: Sorting neuronal and non-neuronal nuclei for RNA-Seq analysis.** Nuclei from the hippocampal CA region were subjected to FANS as depicted in Fig 2A. **A.** Representative images showing nuclei that were sorted using the neuronal marker NeuN. Note that no NeuN positive nuclei are detected in the NeuN (-) fraction confirming the purity of the approach. Scale bar: 50μm **B.** Gating strategy for NeuN (+) and NeuN (-) nuclei sorting. C. RNA-sequencing (n=2/group) was performed from NeuN (+) and NeuN (-) nuclei and a differential expression analysis was performed. Heat map shows 836 genes specifically enriched in NeuN (+) nuclei when compared to NeuN (-) nuclei. The criteria to select those genes were: adjusted *p*-value < 0.01, basemean > 150, fold change > 5. **D.** GO-term analysis showing that the top 10 enriched biological processes and molecular functions for the 836 genes enriched in NeuN (+) nuclei all represent specific neuronal processes. **E.** Normalized expression values obtained from the RNA-seq experiment showing the expression of selected genes known to be enriched in neurons. F. Normalized expression values of genes that are known to be enriched in non-neuronal cells including glia cells. Error bars indicate SEM.

**Expanded view Fig. 6. Confirmation of RNA-seq results via qPCR.** Randomly selected genes that were either significantly up- (5 genes) or downregulated (5 genes) in the RNA-seq dataset comparing control vs *Setd1b* cKO mice were tested via qPCR for validation. The graph shows the corresponding log2 (fold changes) in each experiment. The RNA-seq and qPCR results show highly significant correlation.

**Expanded view Fig. 7. Genes downregulated in *Setd1b* cKO mice also exhibit reduced H3K9ac and H3K27ac levels. A.** Profile plots showing H3K9ac and H3K27ac levels at genes with decreased H3K4me3 in *Setd1b* cKO. **B.** Profile plots showing H3K9ac and H3K27ac levels at downregulated (upper panels) and an equally sized random set of unchanged genes (lower panels) in *Setd1b* cKO. Bar graphs on the right of each profile plot show quantifications of the enrichment of the histone mark at corresponding genomic regions away from TSS (Student t-test: * *p*-value < 0.05, ** *p*-value < 0.01, *** *p*-value < 0.001).

**Expanded view Fig. 8:** Genome browser views showing the 4 analyzed four histone modifications and the gene expression pattern for the *Npy2r* and *Klk8* genes that were previously shown to be involved in synaptic plasticity and learning & memory.

**Expanded view Fig. 9: Comparison of gene expression changes in *Setd1b* cKO mice detected from cell type specific and bulk tissue RNA-seq. A.** Venn diagram showing the overlap of genes downregulated in *Setd1b* cKO mice detected via RNA-seq from neuronal nuclei or bulk hippocampal CA tissue. Importantly, the majority of the downregulated genes detected in bulk tissue RNA-seq are also detected in RNA-seq from neuronal nuclei (281 out 486). Please note that more downregulated genes are detected when neuronal nuclei are analyzed suggesting that some of the differences may be masked by cell type heterogeneity when bulk tissue is analyzed. 205 genes are exclusively detected in the bulk tissue analysis. This might be explained by the fact that especially the cytoplasmatic and synaptic transcriptome is subjected to dynamic changes in post-transcriptional modifications such a RNA-methylation and the regulation by microRNAs, that may even originate from other cells in a paracine manner {Epple, 2021}. **B-D.** The view that both transcriptomes reflect relevant changes due to the loss of *Setd1b* is supported by GO analysis revealing similar pathways that are linked to neuronal plasticity, and the fact that within the TOP 30, 8 GO-terms are identical, amongst then for example “positive regulation of ERK1 and ERK2 cascade”, “memory” or “positive regulation of axon extension”.

**Expanded view Fig. 10: Comparison of the genes downregulated in *Setd1a* heterozygous mutant and *Setd1b* cKO mice.** Venn diagram showing genes downregulated in *Setd1a* heterozygous mutant vs *Setd1b* cKO mice. Please note that genes affected in the different mutant mice are very different. The data from *Setd1a* heterozygous mutant mice stems from a recent publication by Mukai et al., 2019 (PMID:31606247). Care has to be taken since in that study cortical tissue from heterozygous mice constitutively lacking *Setd1a* was analyzed, while our data stems from the hippocampus of conditional knockout mice.

**Expanded view Fig. 11:** Volcano plots showing differential gene expression in the three KMT cKO mice (data from bulk RNA-sequencing results is compared here) together with the overlay of neuron-specific genes. In accordance with our previous observations, although the total number of downregulated genes in each KMT cKO are comparable, more neuron-specific genes are affected in *Setd1b* cKO, followed by *Kmt2a* cKO and *Kmt2b* cKO.

**Expanded view Fig. 12: A.** Profile plot representing, in wild type mice, the average H3K4me3 distribution around TSS of all genes that display decreased H3K4me3 in *Setd1b* cKO, *Kmt2a* cKO or *Kmt2b* cKO neurons. **B.** Bar plots representing the number of genomic regions in distance to the TSS that show significantly decreased H3K4me3 in *Kmt2a* cKO, *Kmt2b* cKO and *Setd1b* cKO neurons binned into 0-2 kb and 0-5 kb upstream or downstream of the TSS. Note that more regions are detected in *Setd1b* cKO neurons when the region more distant to the TSS (0-5bk) is analyzed. **C-E.** Pie charts depicting the genomic distribution of regions with decreased H3K4me3 in *Kmt2a* cKO (C), *Kmt2b* cKO (D) and *Setd1b* cKO (E) neurons.

**Expanded view Fig. 13: Functional categories affected in *Kmt2a* and *Kmt2b* cKO mice. A.** Heat map showing functional GO category analysis for genes downregulated in *Kmt2a* cKO mice. Enrichment of the same categories is also shown for *Kmt2b* and *Setd1b* cKO mice. **B.** Heat map showing functional GO category analysis for genes downregulated in *Kmt2b* cKO mice. Enrichment of the same categories is also shown for *Kmt2a* and *Setd1b* cKO mice. Please note that the GO categories affected in *Kmt2a* or *Kmt2b* cKO mice differ substantially from those affected in *Setd1b* cKO mice. All analyses are based on comparable RNA-seq data generated from bulk hippocampal CA region.

**Expanded view Fig. 14: Comparison of H3K4me3 changes in *Setd1b* cKO, *Kmt2a* cKO, *Kmt2b* cKO and CK-p25 mice.** Overlap of TSS regions (+/− 2kb) with decreased H3K4me3 in each of the three KMT cKO mice with those decreased in mouse model for Alzheimer’s model (CK-p25 mice) (Gjoneska *et al.*, 2015). The patter of overlapping regions (Setd1b > Kmt2a > Kmt2b) is in agreement with our suggested role of the 3 KMTs in the regulation of neuronal genes important for memory function. A highly significant overlap was only seen in case of *Setd1b* cKO mice (580 out of 997 regions; hypergeometric test: *p*-value = 0). The overlap between *Kmt2a* cKO and CK-p25 is much smaller (195 TSS regions) but still remains significant according to the hypergeometric test (*p*-value = 0.00000292). The overlap between *Kmt2b* cKO and CK-p25 is in turn negligible (85 TSS regions) and is not significant (hypergeometric test: *p*-value = 1). It is important to note that we re-analyzed our ChIP-seq data from *Kmt2a*, *Kmt2b* and *Setd1b* cKO mice together with the data from the CK-p25 mice and selected only genomic regions which showed H3K4me3 signal in all datasets to allow for a reliable comparison. Thus, the total numbers of H3K4me3 regions that differ between conditions are slightly different to our results reported for example in Fig. 4A. This is also due to the fact that Gjoneska et al. analyzed bulk hippocampal tissue, whereas our experiments are based on sorted hippocampal neurons from the CA region. Nevertheless, the fact that the overlap is still substantial with *Setd1b* cKO and lower or negligible in the other two KMT cKOs supports the view that *Setd1b* is particularly important for regulating genes important for synaptic plasticity and memory function.

**Expanded view Fig. 15. Comparative analysis of learning and single nucleus gene expression of *Kmt2a*, *Kmt2b* and *Setd1b*. A.** Comparison of the escape latency in the 3 different KMT cKO mice during 10 days of training. Although we applied the same experimental setting, the experiments were performed at different time points ((Kerimoglu *et al.*, 2013) (Kerimoglu *et al.*, 2017), and this study). Thus, for comparison we normalized the data to the corresponding control group. In this plot an increase in the fold change of the normalized escape latency depicts the difference to the corresponding control group. Hence, a higher fold change oft he normalized escape latency indicates a greater difference to the corresponding control and thus more severe learning impairment. *Setd1b* cKO (n = 14) vs *Kmt2a* cKO (n = 13): Repeated measures ANOVA, genotype effect: F (1,25) = 16.83, *** *p*-value < 0.001. *Setd1b* cKO (n = 14) vs *Kmt2b* cKO (n = 22): Repeated measures ANOVA, genotype effect: F (1,34) = 70.66, **** *p*-value < 0.0001. **B.** UMAP plot showing the data from 15661 nuclei. Upper panel shows the different clusters and their corresponding numbers. The lower panel shows the same UMPA plot colored for different groups of cells that are further analyzed in (C) and (D). **C.** Violin plot showing the expression of the 6 KMTs as well as marker genes for the different hippocampal cell types according the UMAP shown in lower panel B. Note that *Kmt2a* and *Kmt2c* are the highest expressed KMTs across cell types, while the other KMTs shown low to moderate expression. **D.** Violin plot showing the expression of the six KMTs and corresponding marker genes within the different cluster representing inhibitory and excitatory neurons. Note that there is no obvious difference of KMT expression between cell types. In agreement with the data shown in panel (C), *Kmt2a* and *Kmt2c* exhibit the highest expression levels. **D.** Bar charts showing the total number of reads/cell that are positive for the corresponding KMT. **E.** Left panel shows representative images of RNAscope performed for *Kmt2a*, *Kmt2b* and *Setd1b*. Right panel shows a bar chart quantifying the dots/cell (n = 1500 cells from 2 mice) indicative of the corresponding expression level. *Kmt2a* expression was significantly higher when compared to *Kmt2b* or *Setd1b* (*** *p*-value < 0,0001, Student t-test).

