## Supplementary figures and images for "Postnatal SETD1B is essential for learning and the regulation of neuronal plasticity genes"

### Expanded View Figure 1

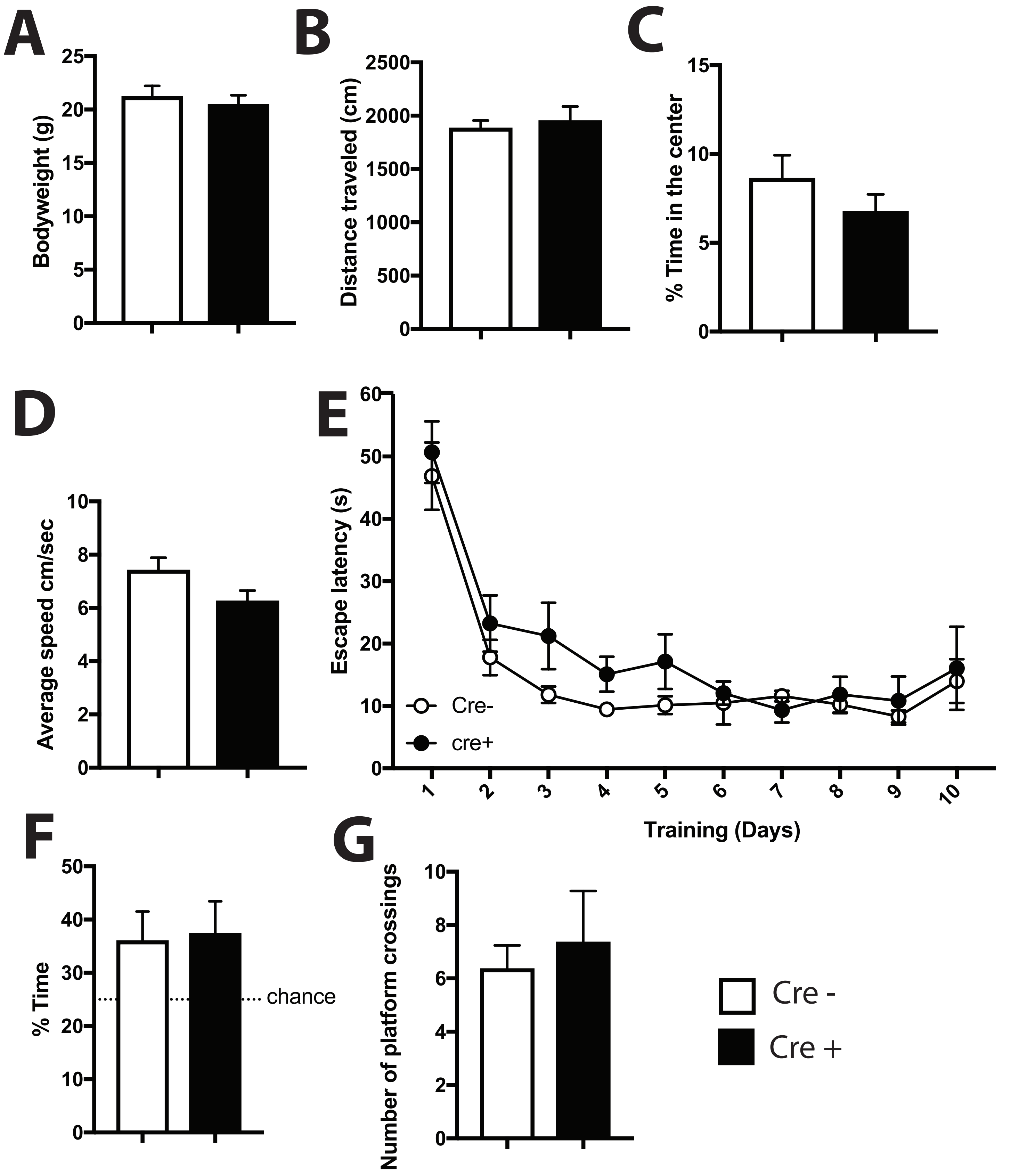

### Expanded View Figure 2

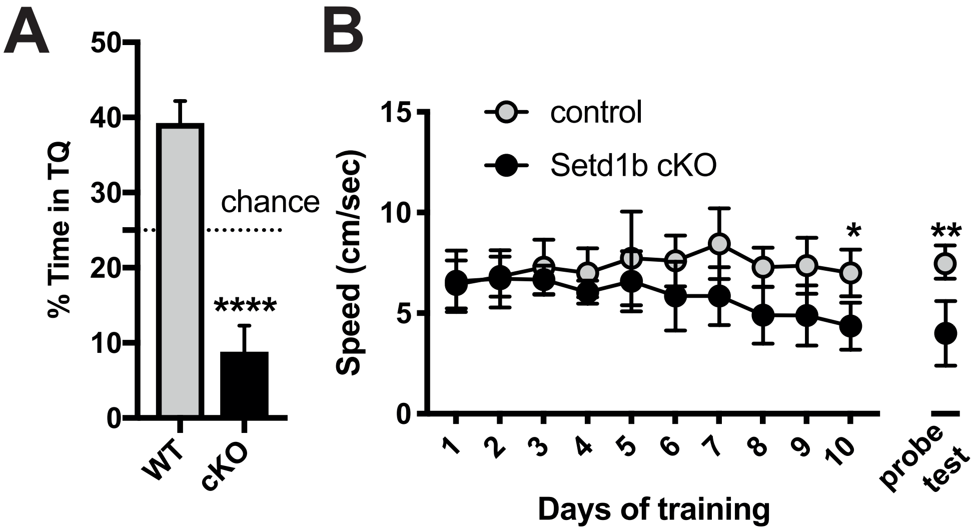

### Expanded View Figure 3

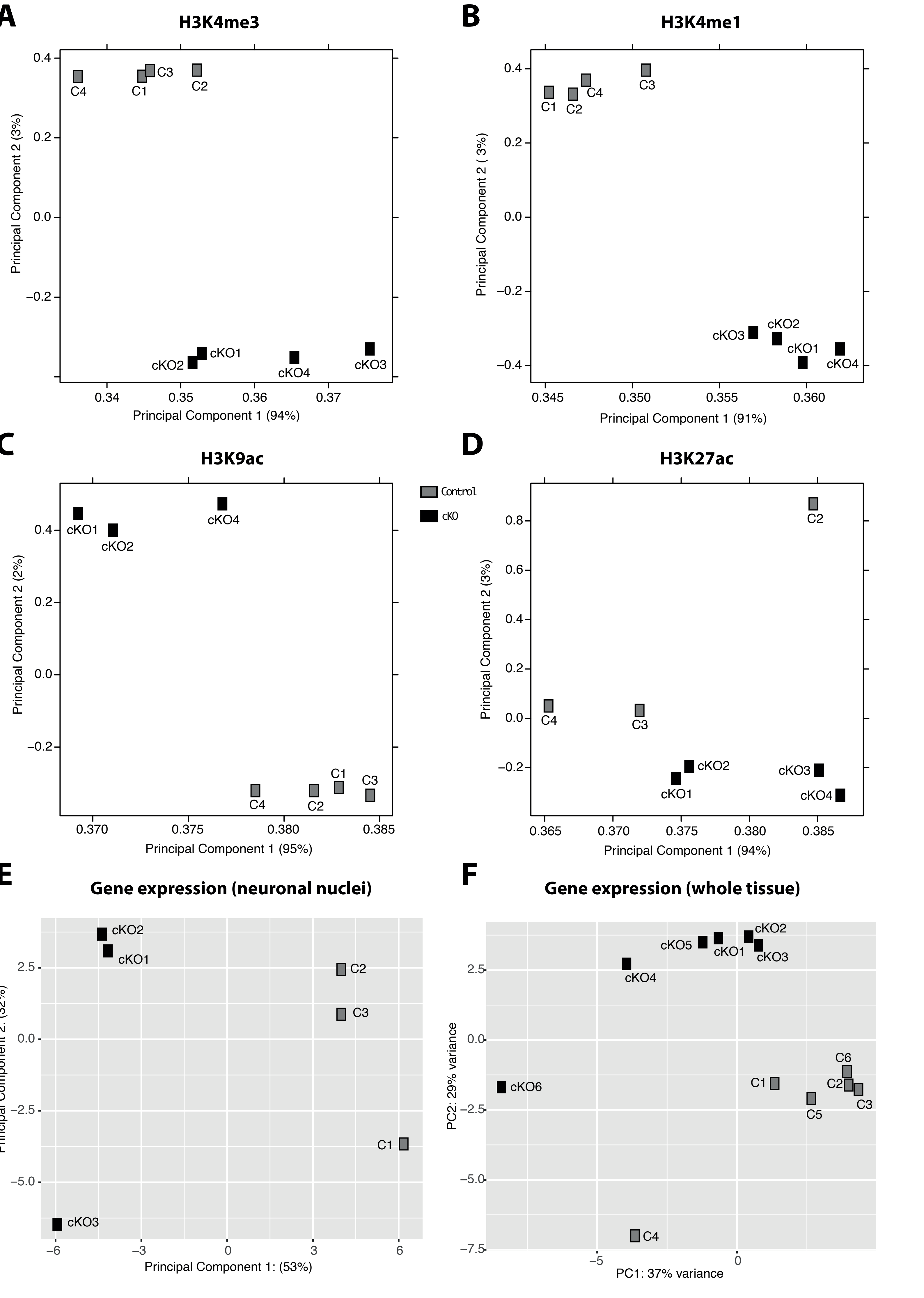

### Expanded View Figure 4

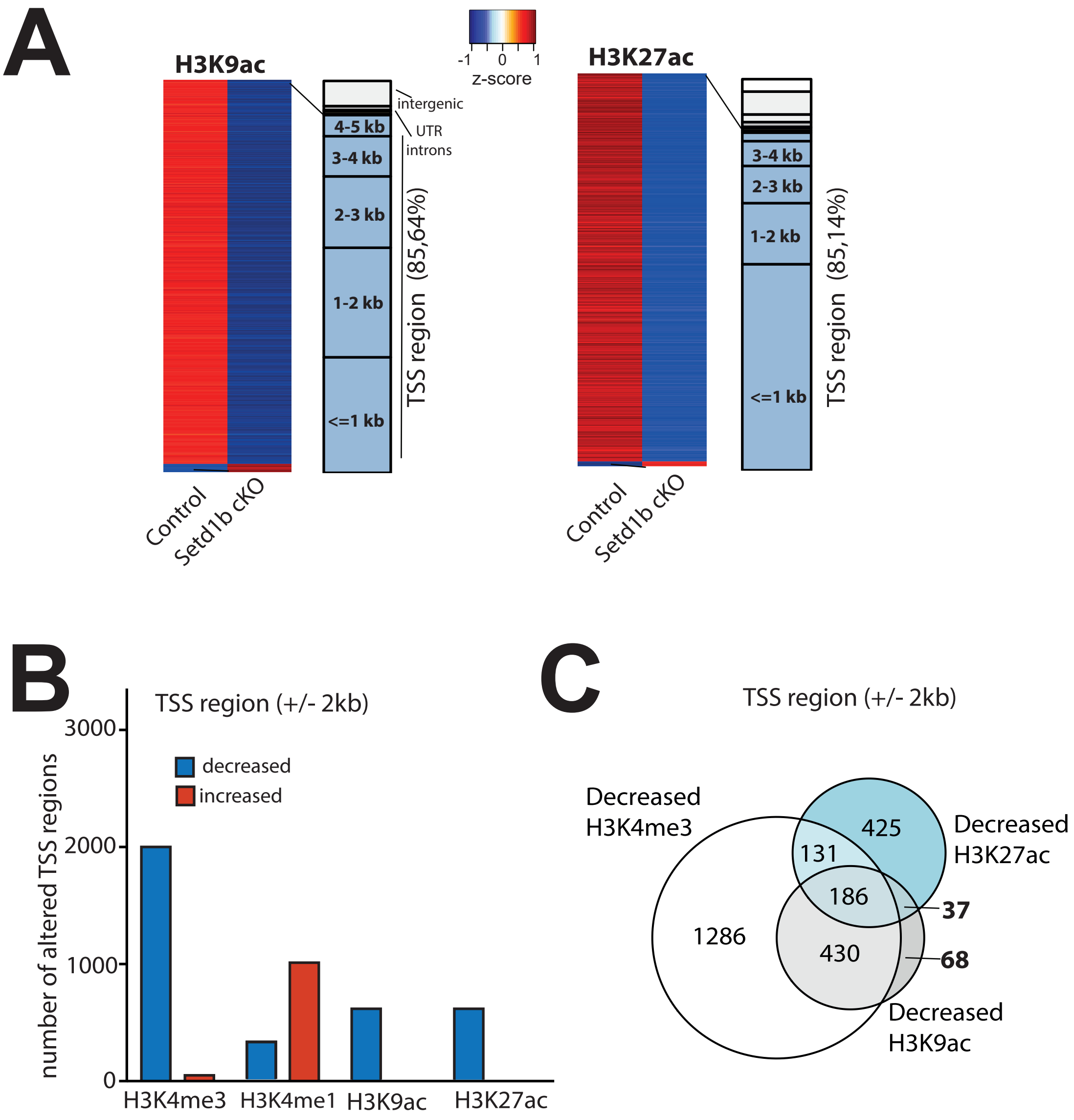

### Expanded View Figure 5

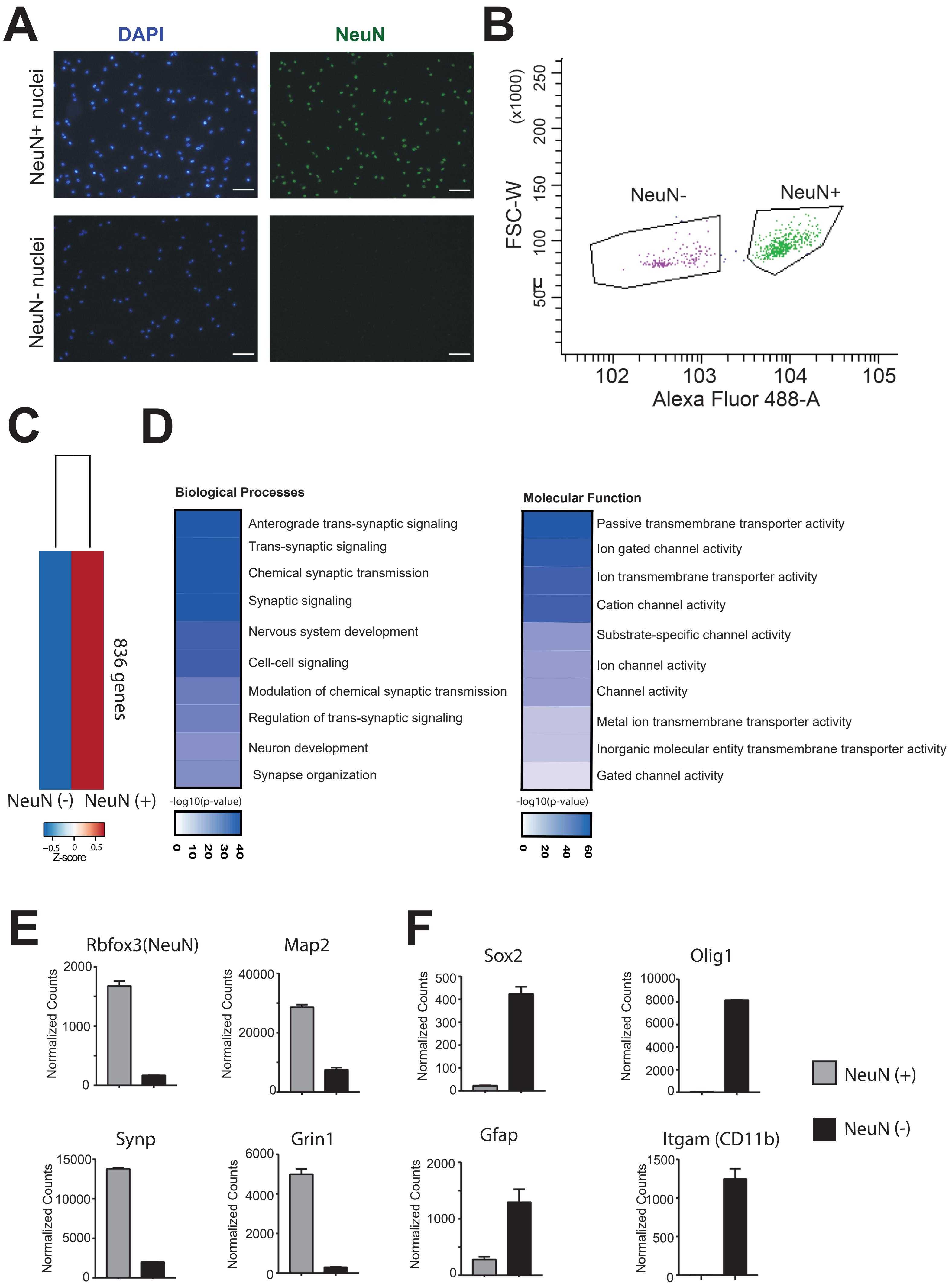

### Expanded View Figure 6

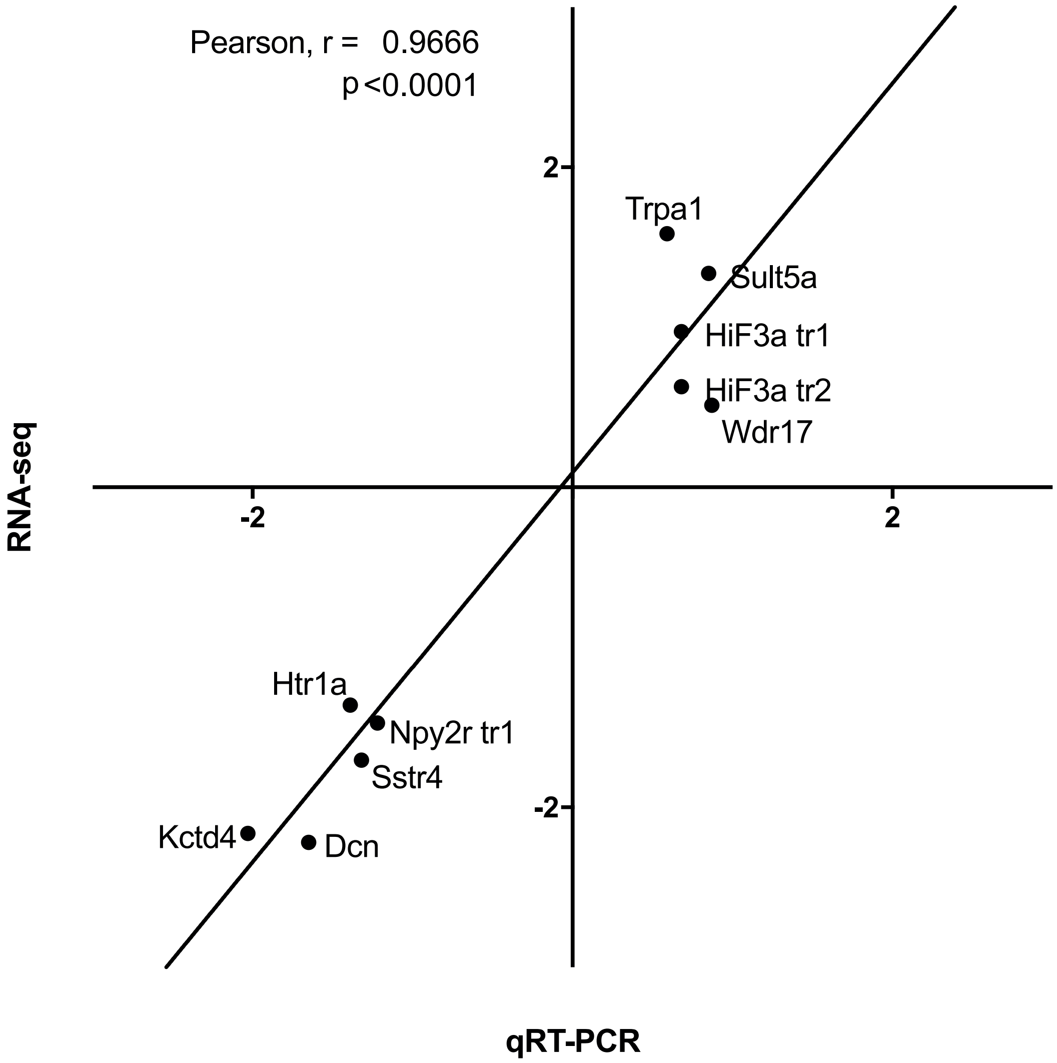

### Expanded View Figure 7

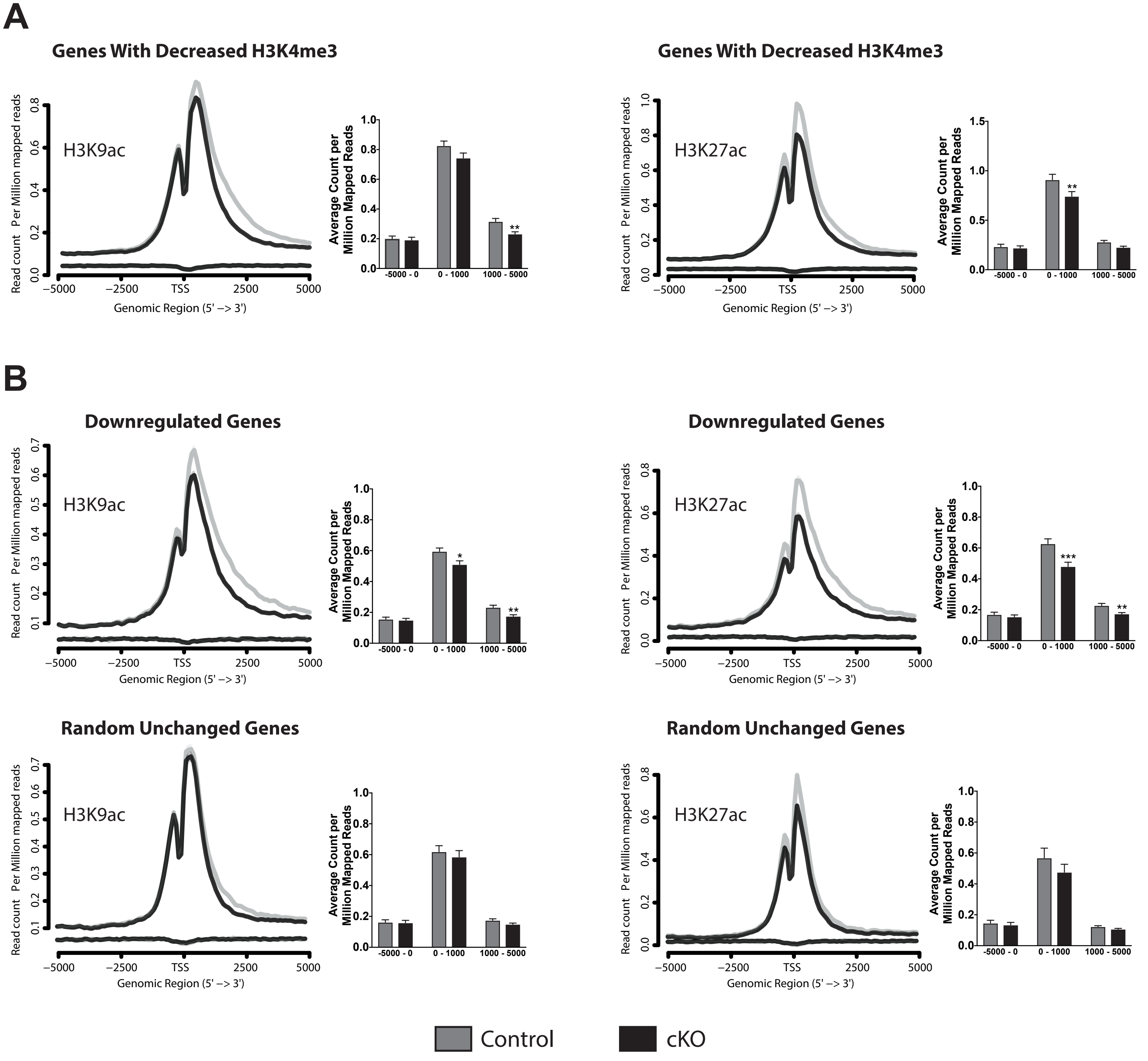

### Expanded View Figure 8

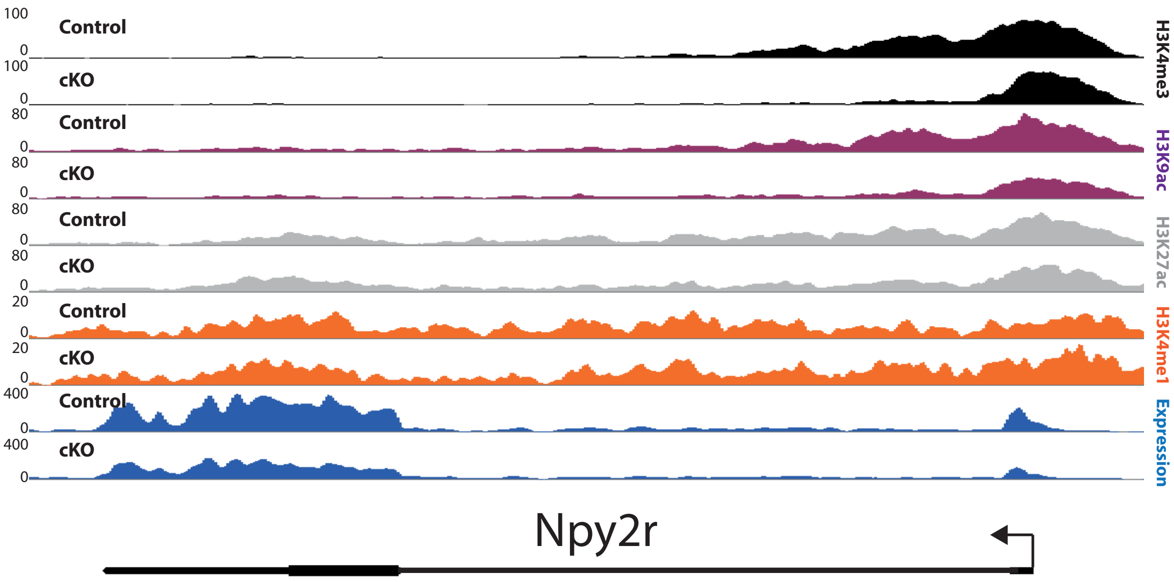

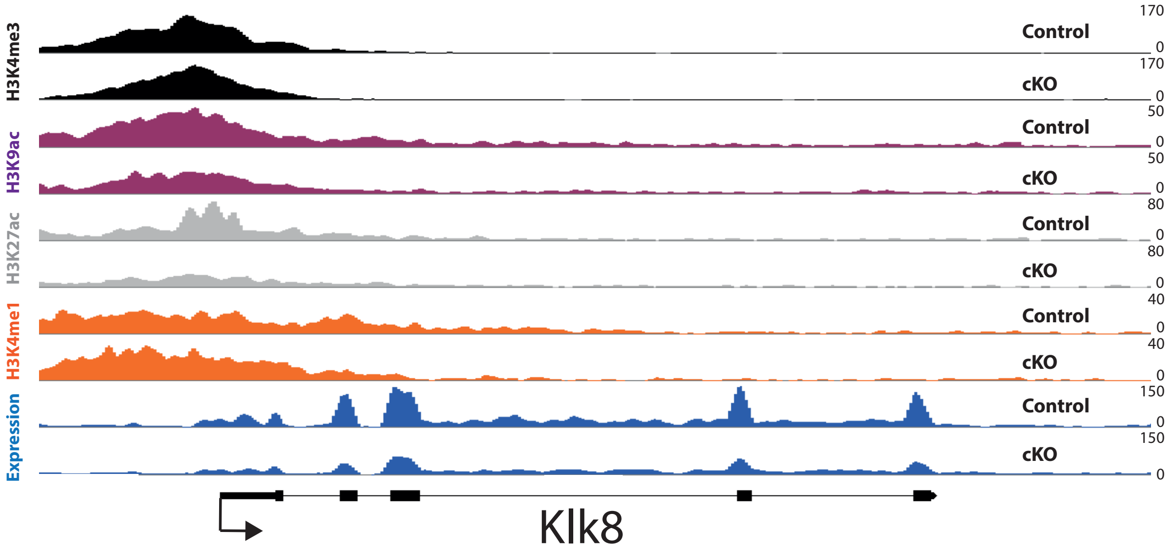

### Expanded View Figure 9

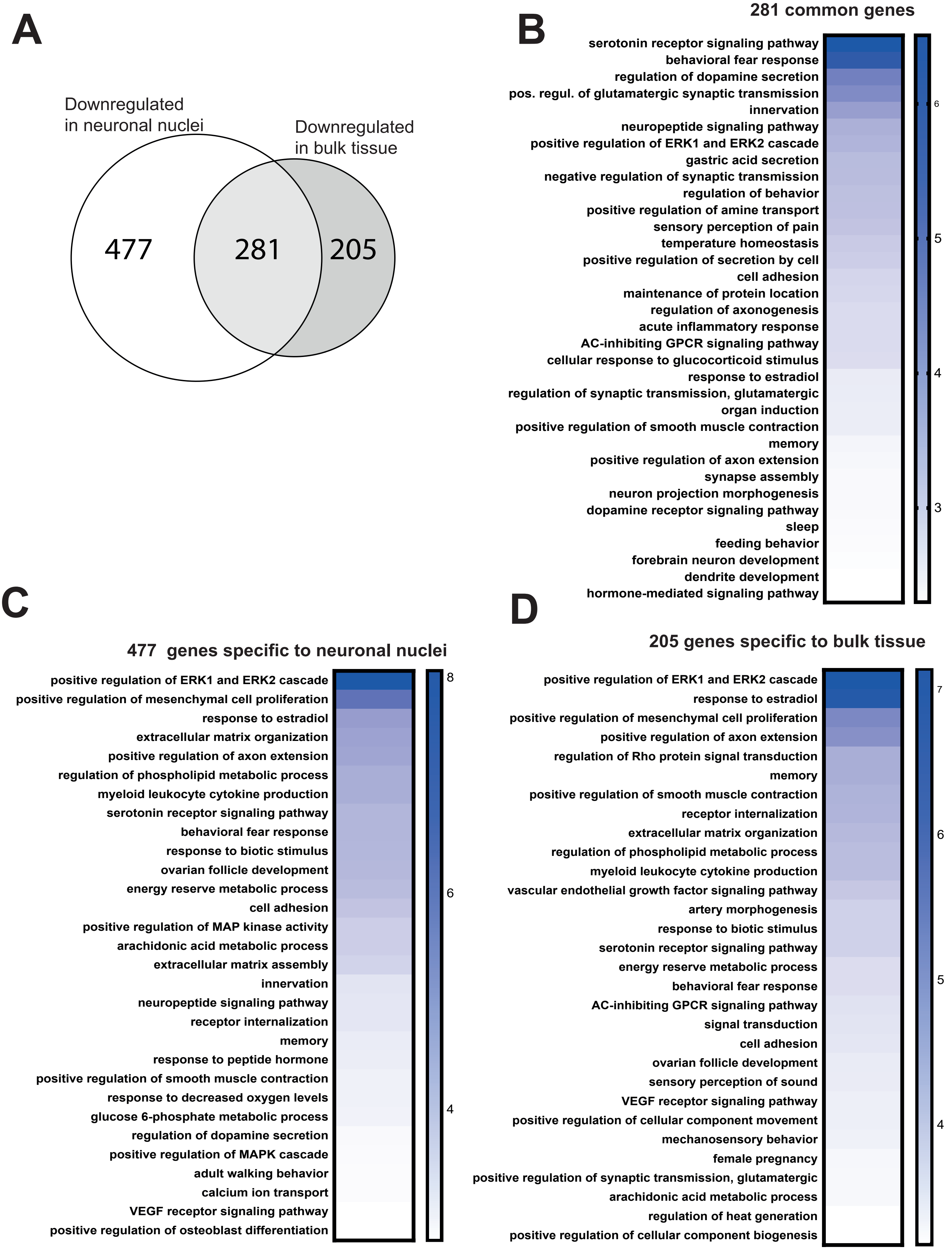

### Expanded View Figure 10

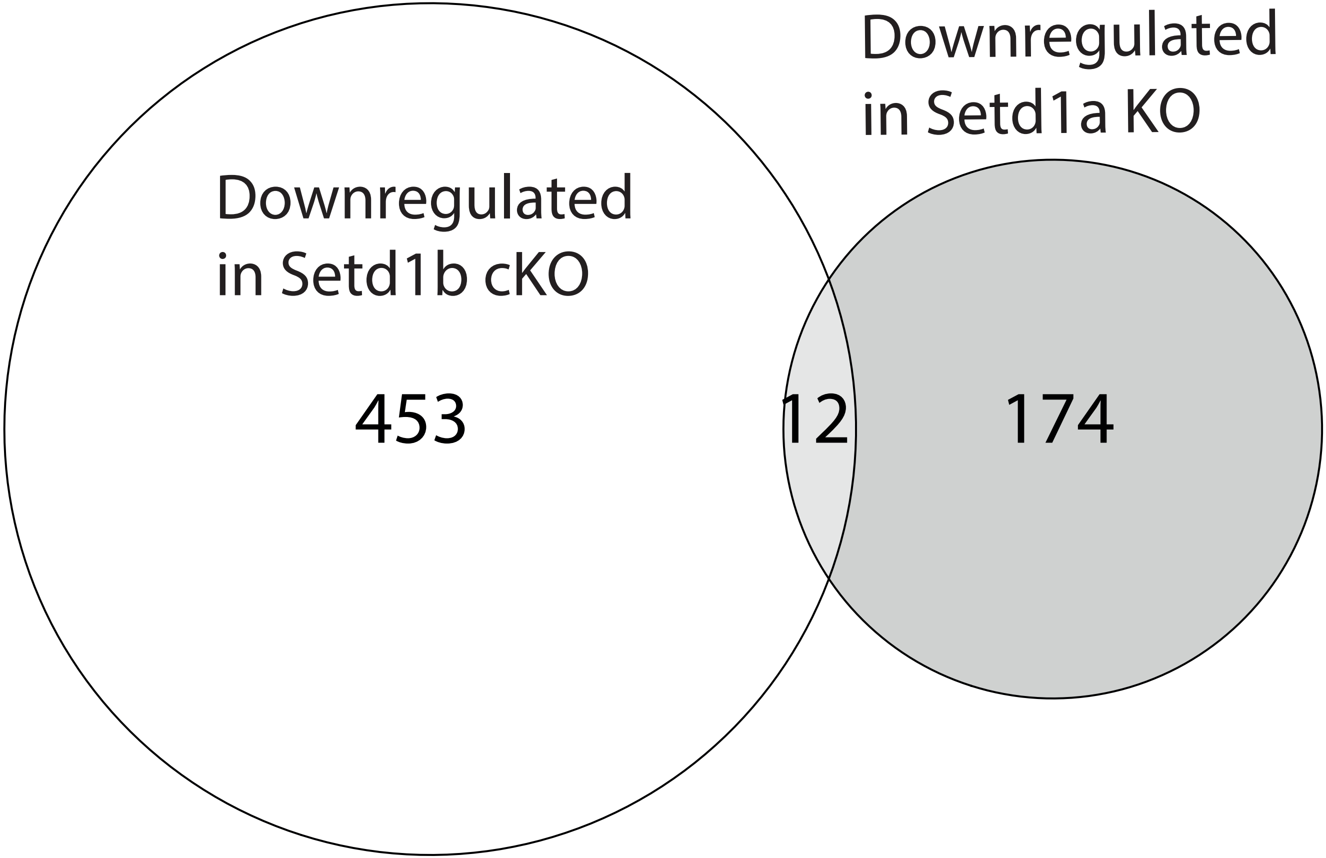

### Expanded View Figure 11

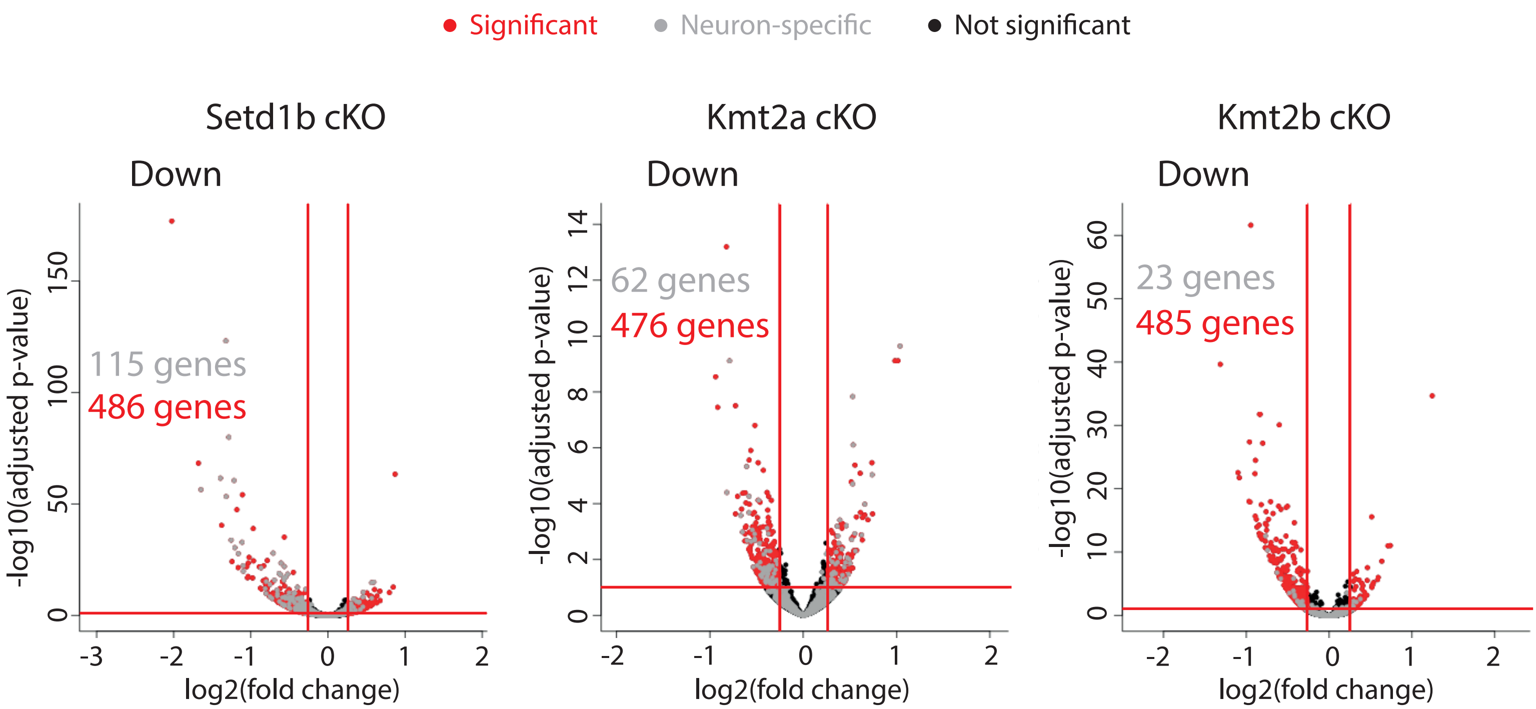

### Expanded View Figure 12

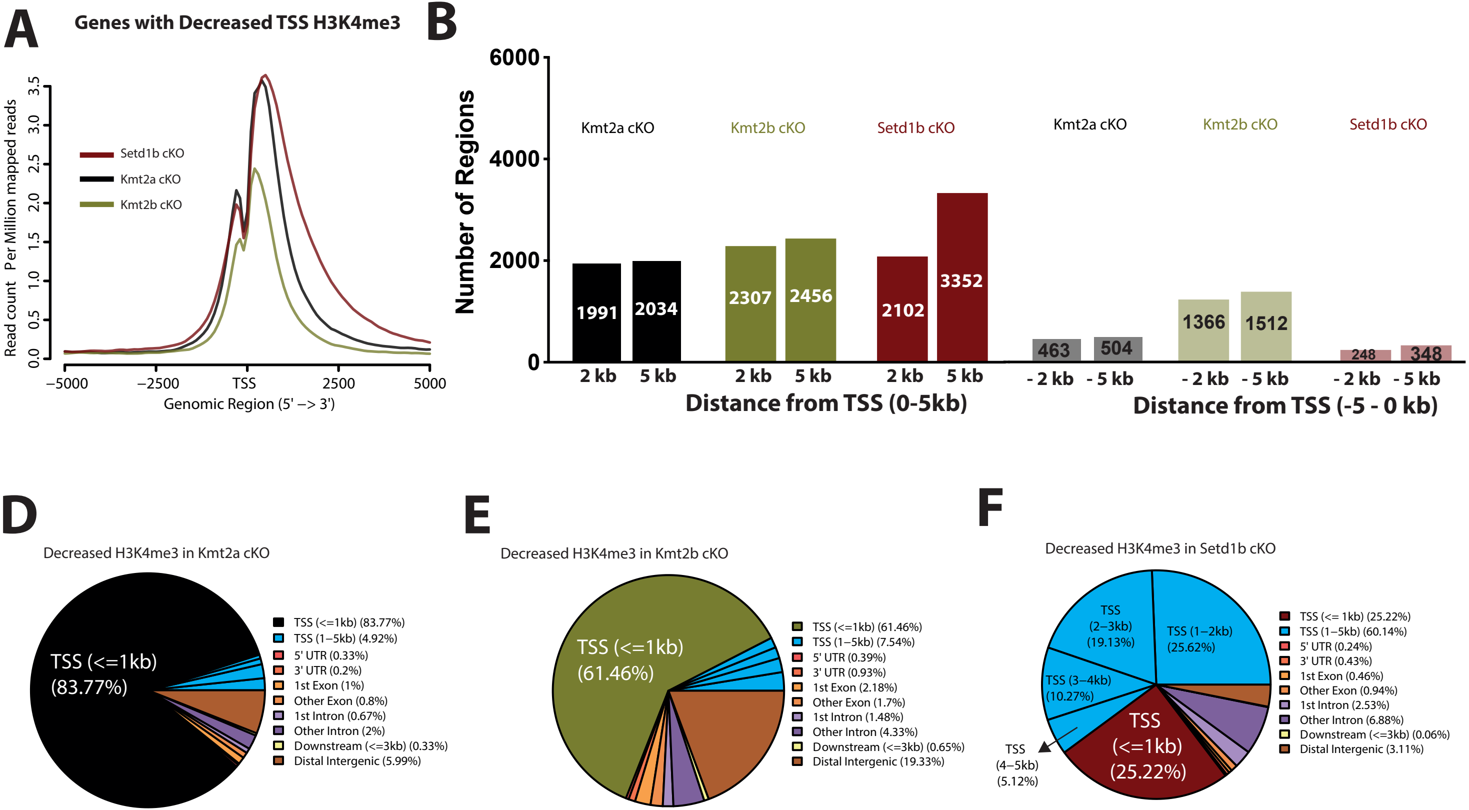

### Expanded View Figure 14

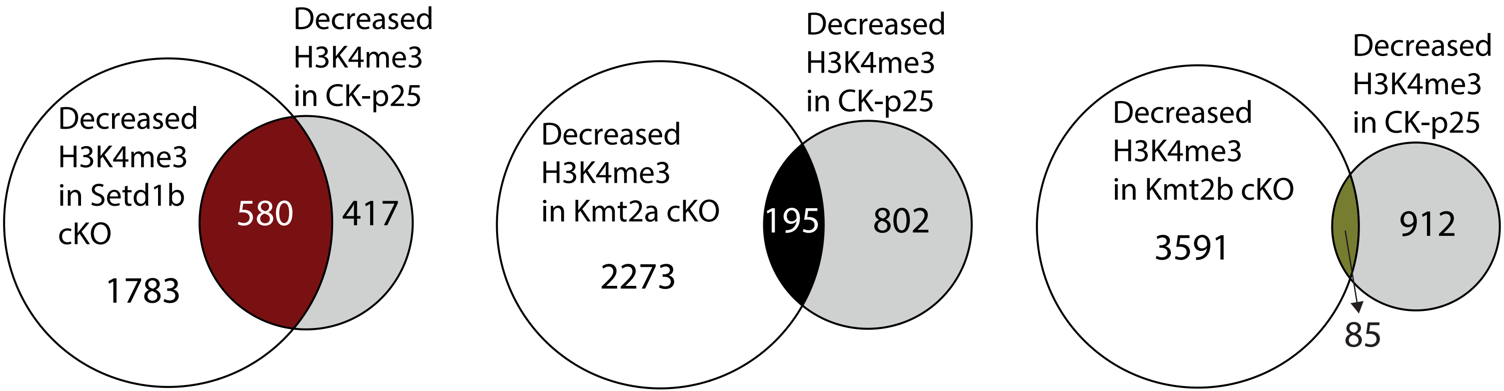

### Expanded View Figure 15

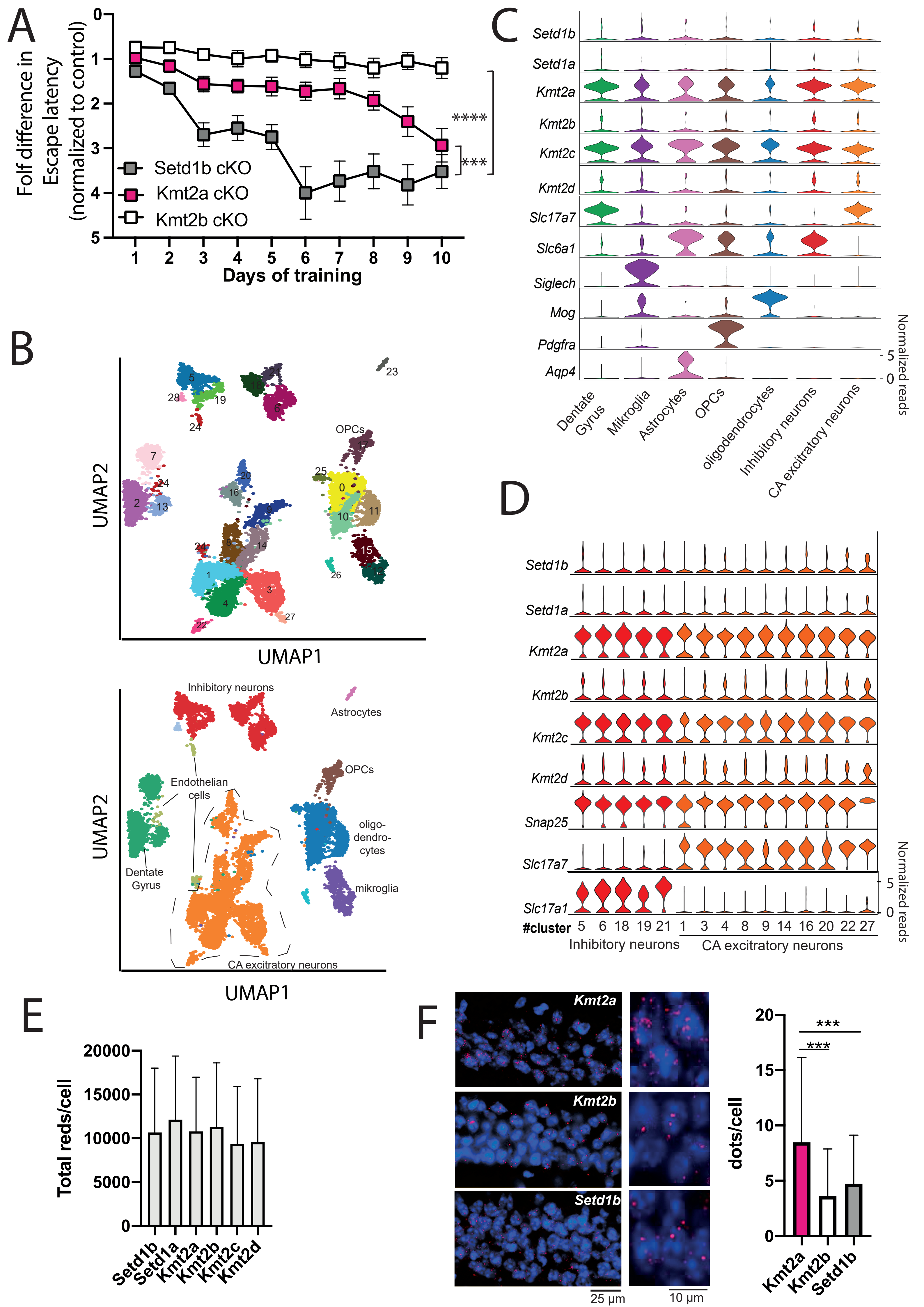
