## Supplementary material for "Postnatal SETD1B is essential for learning and the regulation of neuronal plasticity genes": Expanded View Figure 13

**A**

### Downregulated Genes with Decreased H3K4me3 in cKO

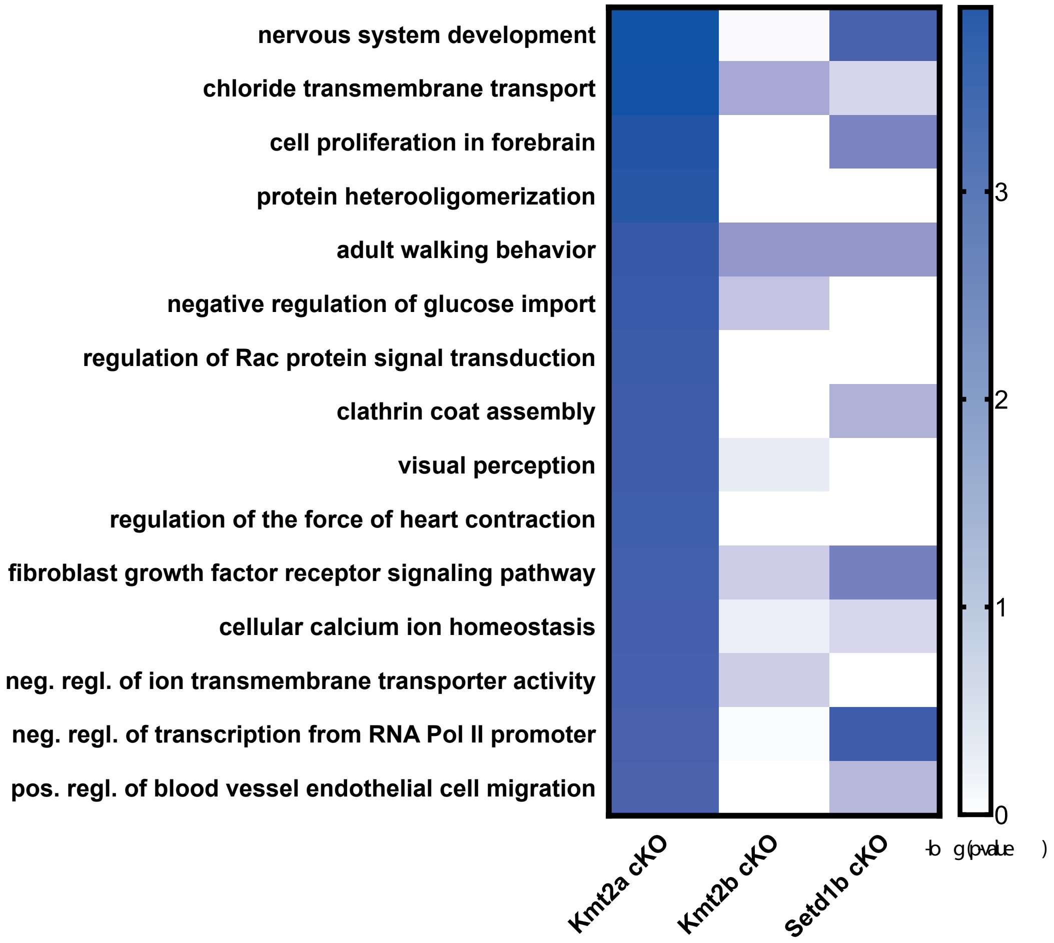


**B**

### Downregulated Genes with Decreased H3K4me3 in cKO

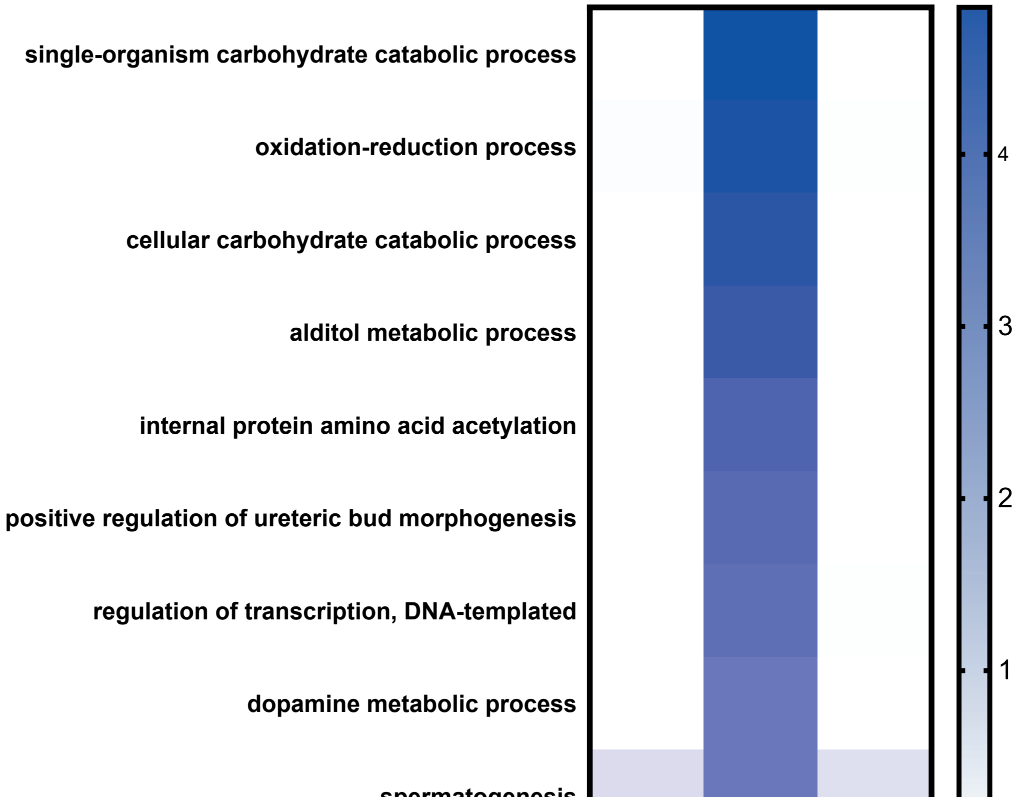
